# Characterization and description of *Clostridium filamentum* ETTB, a novel gut bacterium with TLR5 modulating properties

**DOI:** 10.1101/2025.02.28.640726

**Authors:** Kassem Makki, Maria Elena Martino, Antton Alberdi, Ostaizka Aizpurua, Andrea Quagliariello, Lisa Olsson, Sara Clasen, Johanna Jönsson, Harald Brolin, Chinmay Dwibedi, Gaohua Yang, Chiara Favero, Per-Olof Bergh, Pamela Schnupf, Ruth Ley, Muhammad-Tanweer Khan, Valentina Tremaroli, Fredrik Bäckhed

**Affiliations:** The Wallenberg Laboratory, Department of Molecular and Clinical Medicine, Sahlgrenska Academy, University of Gothenburg, Gothenburg, Sweden; Department of Comparative Biomedicine and Food Science, University of Padova, Padova, Italy; Center for Evolutionary Hologenomics, Globe Institute, University of Copenhagen, Copenhagen, Denmark; Department of Microbiome Science, Max Planck Institute for Biology, Tübingen, Germany; Institut Necker Enfants Malades, Inserm U1151, CNRS UMR8253, Laboratory of Host-Microbiota Interactions, Paris, France; Region Västra Götaland, Sahlgrenska University Hospital, Department of Clinical Physiology, 41345 Gothenburg, Sweden

**Keywords:** Clostridium, TLR5, metagenomes

## Abstract

The *Clostridium* genus is highly heterogeneous, encompassing numerous species and strains, many of which remain to be isolated and characterized to better understand their relationship to host physiology. This study aimed to isolate and characterize novel bacterial species within the Clostridium genus and explore their potential links to host health. Under microaerophilic conditions, we isolated and characterized three bacterial isolates belonging to a new anaerobic *Clostridium* species, designating *Clostridium* sp. DSM 115107 (*Clostridium filamentum* ETTB3) as the type strain. *C. filamentum* ETTB isolates are rod- to filament-shaped, Gram-positive bacteria and exhibit poor growth when cultured on rich media such as LYBHI. Genome sequencing and phylogenetic analysis revealed that *C. filamentum* ETTB belongs to the *Clostridium* genus and clusters closely with *Clostridium saudiense* JCC. Interestingly, *C. filamentum* ETTB has a significantly smaller genome compared to *C. saudiense* JCC containing a reduced repertoire of genes involved in carbohydrate degradation and amino acid synthesis and a larger number of genes related to cell motility, including an additional copy of the *fliC* gene. Unlike *C. saudiense*, *C. filamentum* ETTB adopted a filamentous morphology when in contact with Caco-2 cells and stimulate the TLR5 pathway in Caco-2 cells. Metagenomics analysis revealed that *C. filamentum* ETTB is present in both industrialized and non-industrialized populations, although the relative abundance varying considerably between and within individuals. Our study identifies a novel bacterial strain adapted for the human gut that has the potential to influence host immune response by activating TLR5 pathway.

## Introduction

The human gut hosts trillions of microbial cells that form an intricate network, regulating host physiology^1^. Despite the advances in knowledge brought by metagenomics, approximately 70% of gut bacteria detected in metagenomics studies lack cultured representatives^2^, which has spurred efforts to isolate and characterize human gut bacteria with the potential to identify novel interactions with the host.

Research has shown that the *Clostridium* genus can regulate several essential biological processes within the host, including gut integrity, immune functions, and glucose metabolism^3–6^. However, the *Clostridium* genus is remarkably diverse, encompassing unidentified, uncultured, or poorly characterized species. This diversity extends not only to different species but also to strains within species, as highlighted by recent studies^7–10^. This heterogeneity includes both beneficial, commensal clostridia as well as pathogenic strains, indicating their potential roles in either maintaining host health or contributing to disease development through various mechanisms. Therefore, the isolation and characterization of new *Clostridium* species hold significant promise for advancing our understanding of bacterial interactions with host cells.

In this study, we characterize three novel isolates obtained from the stools of a healthy individual, representing a previously unidentified *Clostridium* species. These bacteria were characterized at the morphological and metabolic levels, and *in silico*, *in vitro*, and metagenomic studies were used to explore functions relevant to their interaction with the host. Based on morphological characteristics, we propose the name: *Clostridium filamentum* ETTB, with *Clostridium* sp. DSM 115107 (*C. filamentum* ETTB3) as the type strain.

## Results

### Phenotypical and genetic characterizations of *Clostridium filamentum* ETTB

To isolate bacteria present in the human gut, we collected a fecal sample from a healthy volunteer and cultured it in a bioreactor designed to mimic the environment of the human colonic gut. The bioreactor contained one anaerobic chamber and one oxygenated chamber, separated by an agar septum coated with porcine gastric mucin allowing oxygen diffusion towards the anaerobic chamber (Supplementary Figure 1A). Several colonies were retrieved from the septum, including three filamentous bacteria showing similar morphological characteristics (Figure 1A and 1B). These isolates were designated as *Clostridium filamentum* ETTB1, ETTB2, and ETTB3, with ETTB3 as the type strain for the novel species *C. filamentum* deposited at DSMZ with name *Clostridium* sp. DSM 115107 and CCUG with the name CCUG 77667. Of note, analysis of the 16S rRNA gene revealed 98% homology with the gene sequence of *Clostridium saudiense* JCC (Supplementary information 1). Accordingly, *C. saudiense* JCC was used as a comparator to the new isolates.

**Figure 1:**
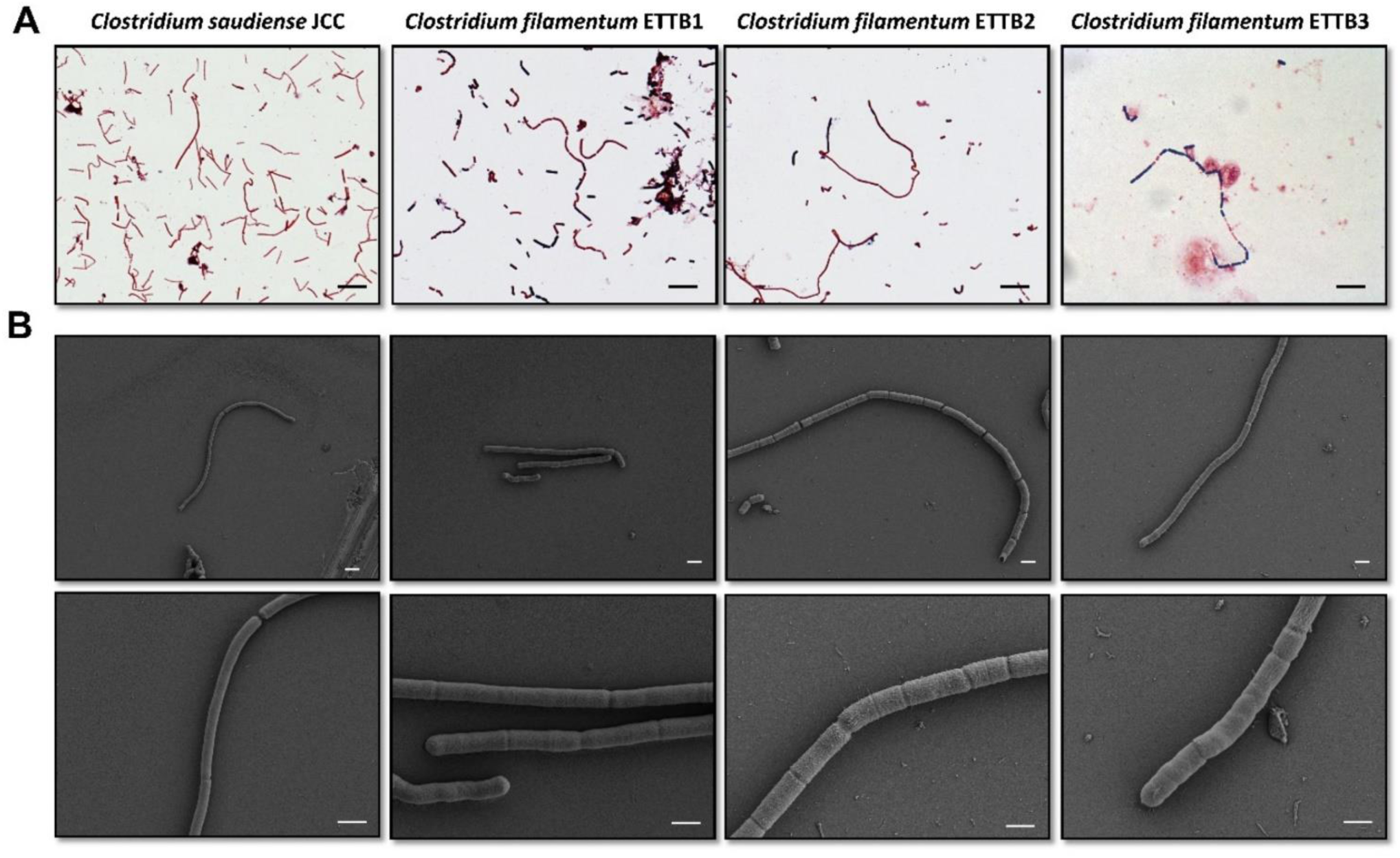
Morphological characterization of *C. saudiense* JCC and *C. filamentum* ETTB1, ETTB2 and ETTB3. A. Gram staining of overnight cultures grown in LYBHI broth. B. Scanning electron microscopy showing the morphology of *C. saudiense* JCC and *C. filamentum* isolates. For acquiring SEM pictures, electron high tension (EHT) of 2.00 KV and magnifications of 7.00 KX (upper panel B) and 25.00 KX (bottom panel B) were used. Scale bars represent 5 µm length for panel A and 2 µm length for panel B.

*C. filamentum* ETTB isolates exhibit Gram-variable and polymorphic characteristics (Figure 1A and Supplementary Figure 1B) and are highly sensitive to ambient oxygen, as evidenced by the lack of growth in aerobic conditions. No growth was observed for these isolates when exposed for 10 minutes to ambient air or in microaerophilic conditions (Table 1). When cultured on LYBHI agar plates for 24 hours, the colonies appeared relatively small (approximately 0.5-1 mm), raised, with silver colour, and circular to irregular borders. A marked difference in morphology was observed between *C. saudiense* JCC and *C. filamentum* ETTB isolates (Figure 1A and Supplementary Figure 1B). Both species possess rod to filamentous-shaped cells (Figure 1A and 1B). However, scanning electron microscopy imaging revealed differences in segmentation patterns and distinct outer cell envelopes for *C. saudiense* JCC and *C. filamentum* ETTB isolates (Figure 1B). The polar lipid profiles also markedly differed between the two species, with the cell wall of *C. saudiense* JCC containing a larger range of lipids mainly glycolipids and aminoglycolipids compared to *C. filamentum* ETTB isolates (Supplementary Figure 1C).

**Table 1.**

| Growth in |  | *C. saudiense*<br>JCC | *C. filamentum*<br>ETTB1 | *C. filamentum*<br>ETTB2 | *C. filamentum*<br>ETTB3 |
| --- | --- | --- | --- | --- | --- |
| Temperature (°C) | 25 | + | Poor | Poor | Poor |
|  | 37 | + | + | + | + |
|  | 42 | + | + | + | + |
| Conc. NaCl (%) | 5 | - | - | - | - |
|  | 1 | + | Poor | Poor | Poor |
|  | 0.2 | + | + | + | + |
| Bovine bile acids (%) | 0.5 | + | + | + | + |
|  | 0.1 | + | + | + | + |
| Motility | 0.5% agar (LyBHI) | + | + | + | + |
| Different pH | 5 | - | - | - | - |
|  | 6 | Poor | Poor | Poor | Poor |
|  | 7 | + | + | + | + |
|  | 8 | + | + | + | + |
|  | 9 | + | Poor | Poor | Poor |
| Oxygen levels | Microaerophilic (5-10%) | - | - | - | - |
|  | Strict anaerobic | + | + | + | + |
| Media | Anaerobic BHI | + | + | + | + |
|  | Columbia Blood Agar | + | + | + | + |
|  | 2YT | + | Poor | Poor | Poor |
|  | YCFAG | + | - | - | - |
|  | Luria-Bertani Broth | Poor | Poor | Poor | Poor |

### Genome comparison of *Clostridium filamentum* ETTB with *Clostridium saudiense* JCC

We performed genome sequencing and confirmed that the three new isolates ETTB1, ETTB2 and ETTB3 belonged to the same bacterial species, as evidenced by their genome size, percentage of GC content, and average nucleotide identity (ANI) of 99.98% between the *C. filamentum* ETTB isolates (Table 2). As shown by the 16S rRNA gene analysis, *C. saudiense* JCC was the most closely related species to *C. filamentum* ETTB (Figure 2A). These observations were further confirmed using Protologger – automated tool for description of novel bacteria^11^. Also, percentage of conserved proteins (POCP) analysis indicated that *C. filamentum* ETTB isolates belong to *Clostridium* genus with POCPu values of 53.88%, 53.87%, 53.86% for *C. filamentum* ETTB1, ETTB2 and ETTB3, respectively.

**Figure 2:**
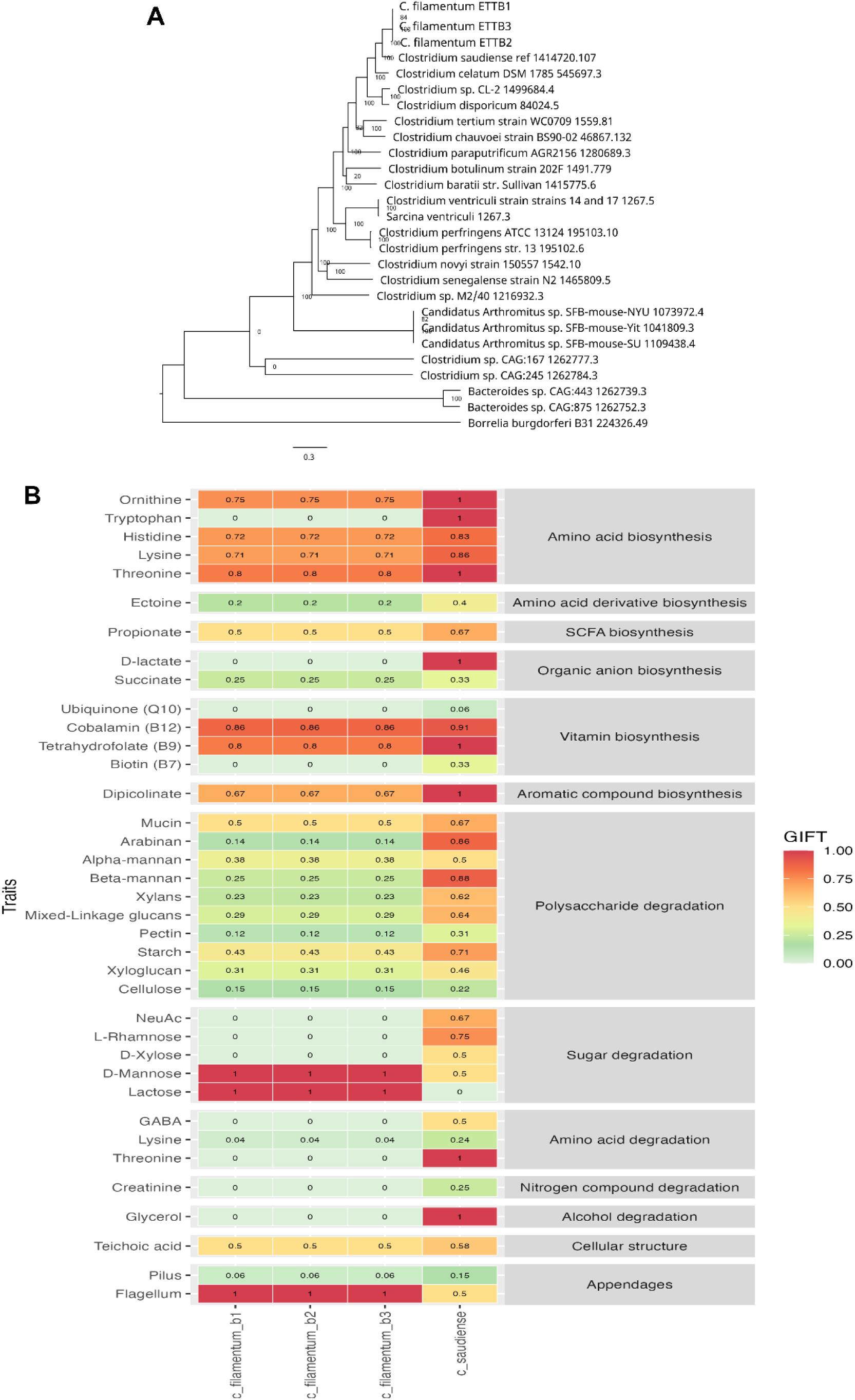
Phylogenetic and genomic analyses for *C. filamentum* ETTB1, ETTB2 and ETTB3 and comparisons with *C. saudiense* JCC. **A.** Phylogenetic tree based on genome sequences showing the relationship of the *C. filamentum* ETTB1, ETTB2 and ETTB3 with other *Clostridium* species. **B.** Pangenome analysis highlighting genomic functional differences with *C. saudiense* JCC.

**Table 2.**

|  |  | *C. saudiense*<br>JCC | *C. filamentum*<br>ETTB1 | *C. filamentum*<br>ETTB2 | *C. filamentum*<br>ETTB3 |
| --- | --- | --- | --- | --- | --- |
| General | GC Content (%) | 28.3 | 27.31 | 27.31 | 27.332 |
|  | Genome Length (bp) | 3 785 077 | 2 820 727 | 2 820 526 | 2 823 133 |
| Annotation | CDS | 3 551 | 2 635 | 2 632 | 2 644 |
|  | Repeat Regions | 118 | 178 | 178 | 179 |
|  | tRNA | 96 | 91 | 91 | 91 |
|  | rRNA | 12 | 11 | 11 | 11 |
|  | partial CDS | 0 | 0 | 0 | 0 |
|  | Hypothetical Protein | 1 133 | 770 | 764 | 780 |
|  | Proteins with functional assignments | 2 418 | 1 865 | 1 868 | 1 864 |
|  | Proteins with EC assignments | 816 | 658 | 660 | 657 |
|  | Proteins with GO assignments | 676 | 551 | 553 | 550 |
|  | Proteins with Pathway assignments | 582 | 475 | 477 | 474 |
| Antibiotic<br>resistance | AMR -CARD |  | 2 | 2 | 2 |
|  | AMR -NDARO |  | 2 | 2 | 2 |
|  | AMR - PATRIC | 28 | 25 | 25 | 25 |
|  | Transporter - TCDB |  | 1 | 1 | 1 |

Interestingly, *C. filamentum* ETTB has a reduced total genome size (Table 2) and the ANI for ETTB1, ETTB2 and ETTB3 with *C. saudiense* JCC was 84.84%, 84.88% and 84.92%, respectively. The phylogenetic tree analysis performed on the whole genome also showed that *C. filamentum* ETTB are closely related to *Clostridium celatum* and *Clostridium disporicum*, but only distantly related to *Clostridium difficile*, *Clostridium tetani* and *Clostridium botulinum,* as well as the segmented filamentous bacteria *Candidatus Arthromitus* present in mice (Figure 2A). *C. saudiense* JCC and *C. filamentum* ETTB isolates also exhibited marked genomic structural differences as highlighted by the linear pangenome analysis (Supplementary figure 2A).

To further elucidate genetic differences between *C. filamentum* ETTB isolates and *C. saudiense* JCC, we conducted a comparative genomic analysis. The clusters of orthologous groups (COG) analysis revealed notable disparities between the two species (Supplementary figure 2B). Specifically, *C. filamentum* ETTB isolates exhibited a lower number of genes involved in carbohydrate transport and metabolism, signal transduction, and defence mechanisms compared to *C. saudiense* JCC (Supplementary figure 2A-B). Conversely, *C. filamentum* ETTB isolates harboured a higher number of genes related to replication, recombination, and repair, as well as translation. Furthermore, *C. filamentum* ETTB isolates possessed a larger number of genes associated with cell motility (Supplementary figure 2B), including genes involved in flagellum formation (Figure 2B). Accordingly, the pangenome analysis revealed marked differences in the presence and completeness of pathways related to *C. filamentum* ETTB ability to synthesize amino and organic acids, as well as its capacity to degrade carbohydrates. Specifically, *C. filamentum* ETTB isolates lacked a complete set of genes involved in the biosynthesis of ornithine, tryptophan, and D-lactate, as well as those responsible for the degradation of carbohydrates and glycerol (Figure 2B).

### *Clostridium filamentum* ETTB are metabolically and functionally different from *Clostridium saudiense* JCC

To highlight metabolic differences between *C. saudiense* JCC and *C. filamentum* ETTB isolates, we assessed the growth of these bacteria on various media (Table 1). *C. filamentum* ETTB strains exhibited a significantly slower growth rate compared to *C. saudiense* JCC in LYBHI (Figure 3A). Furthermore, while both *C. filamentum* ETTB isolates and *C. saudiense* JCC grew anaerobically in BHI and Columbia blood agar, *C. filamentum* ETTB isolates displayed poorer growth on 2YT and failed to grow on YCFAG (Table 1).

**Figure 3:**
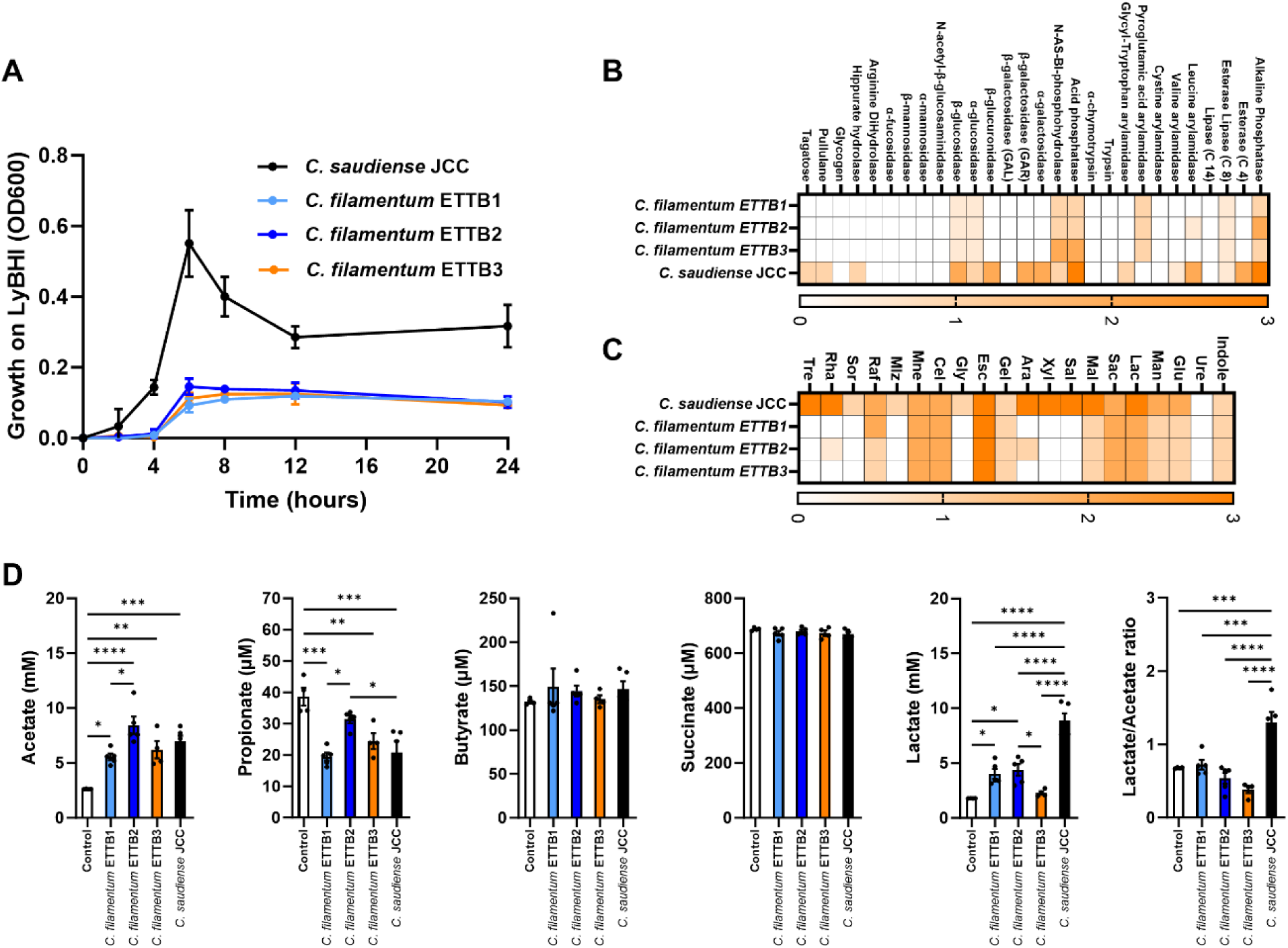
Growth curves and metabolic characterization of *C. filamentum* ETTB and *C. saudiense* JCC. **A.** Growth curves in LYBHI broth for the bacteria incubated anaerobically for 24 hours in a Coy chamber at 37°C. **B** and **C.** Representative heatmaps of 3 independent experiments evaluating enzymatic activities (**B**) and substrate utilization for *C. saudiense* JCC and *C. filamentum* ETTB1, ETTB2 and ETTB3 using API ZYM, API 20A and rapid ID 32 STREP (**C**). **D.** Profiles of short-chain fatty acids produced in LYBHI broth by cultures grown anaerobically for 18 hours in a Coy chamber at 37°C (as described in A). Data are presented as mean ± S.E.M. n= 4-6 independent experiments. One-way ANOVA with post-hoc Tukey’s test: *\*p< 0.05*, *\*\*p< 0.01*, *\*\*\*p< 0.001* and *\*\*\*\*p< 0.0001*.

Compared to *C. saudiense* JCC, *C. filamentum* ETTBs exhibited optimal growth at 37°C, with diminished growth at both 25° and 42°C (Table 1). To evaluate the optimal growth conditions of the isolates across varying levels of acidity, we cultured *C. filamentum* ETTB and *C. saudiense* JCC in LYBHI media at different pH (Table 1). Both strains showed optimal growth at pH 7 and 8, while growth was notably reduced at pH 5 and 6. In contrast, *C. filamentum* ETTB displayed diminished growth compared to *C. saudiense* JCC at pH 9. In terms of salt tolerance, *C. filamentum* ETTB isolates demonstrated growth in the presence of low levels of sodium chloride (0.2% NaCl), but failed to grow in concentrations of 1% and 5% NaCl. Conversely, *C. saudiense* JCC exhibited growth in 0.2% and 1% NaCl, but not in 5% NaCl (Table 1). *C. filamentum* ETTB isolates and *C. saudiense* JCC had similar bile acid tolerance and grew in the presence of 0.1% and 0.5% oxgall (Table 1). Finally, both *C. filamentum* ETTB isolates and *C. saudiense* JCC displayed motility when cultured on 0.5% agar LYBHI plates under anaerobic conditions (Table 1).

Next, we analysed the capabilities of both bacterial species to utilize different substrates. *C. filamentum* ETTBs were unable to degrade complex polysaccharide highlighted by an absence of enzymatic activities for α-GAL, β-Gal and β-Glu degradation (Figure 3B). Also, *C. filamentum* ETTBs were unable to metabolize xylose, arabinose, melezitose, rhamnose and trehalose and metabolized glucose, mannose and maltose less efficiently than *C. saudiense* JCC (Figure 3C). Finally, we characterized short-chain fatty acid, lactate and succinate production in *C. filamentum* ETTB and *C. saudiense* JCC after growth in LYBHI media for 24 hours (Figure 3D). Butyrate and succinate were not produced (Figure 3D). However, *C. filamentum* ETTB1 and ETTB2 and *C. saudiense* JCC but not *C. filamentum* ETTB3 produced lactate. All bacterial isolates produced significant amount of acetate (Figure 3D). These results confirm the reduced capacity of *C. filamentum* ETTB to degrade carbohydrates as also indicated by the genomes analyses (Figure 2B), and showed that *C. filamentum* ETTB produce acetate and small amounts of lactate as end products of fermentation during growth in LYBHI (Figure 3D).

### *C. filamentum* ETTB increase in size and number when grown in co-culture with Caco-2 cells

As *C. filamentum* ETTB were isolated from an agar plug coated with porcine gastric mucin at the interphase between anaerobic and aerobic conditions, we hypothesized that *C. filamentum* ETTB might be able to interact with intestinal epithelial cells. To test this hypothesis, we co-cultured *C. filamentum* ETTB1 and ETTB2 and *C. saudiense* JCC with the colonic Caco-2 cells under anaerobic conditions. It is important to note that *C. filamentum* ETTB3 was not able to grow in a reproducible fashion under these conditions. In contrast, *C. filamentum* ETTB1 and ETTB2 increased their biomass and became more filamentous compared to *C. filamentum* ETTB alone when cultured with Caco-2 cells, while the morphology and biomass of *C. saudiense* JCC did not change when cultured under the same conditions (Figure 4A and Supplementary Figure 3B). Similarly, we did not observe changes in biomass or morphology when *Escherichia coli* or *Lactobacillus reuteri* were incubated with Caco-2 cells (Supplementary Figure 3A-B).

**Figure 4:**
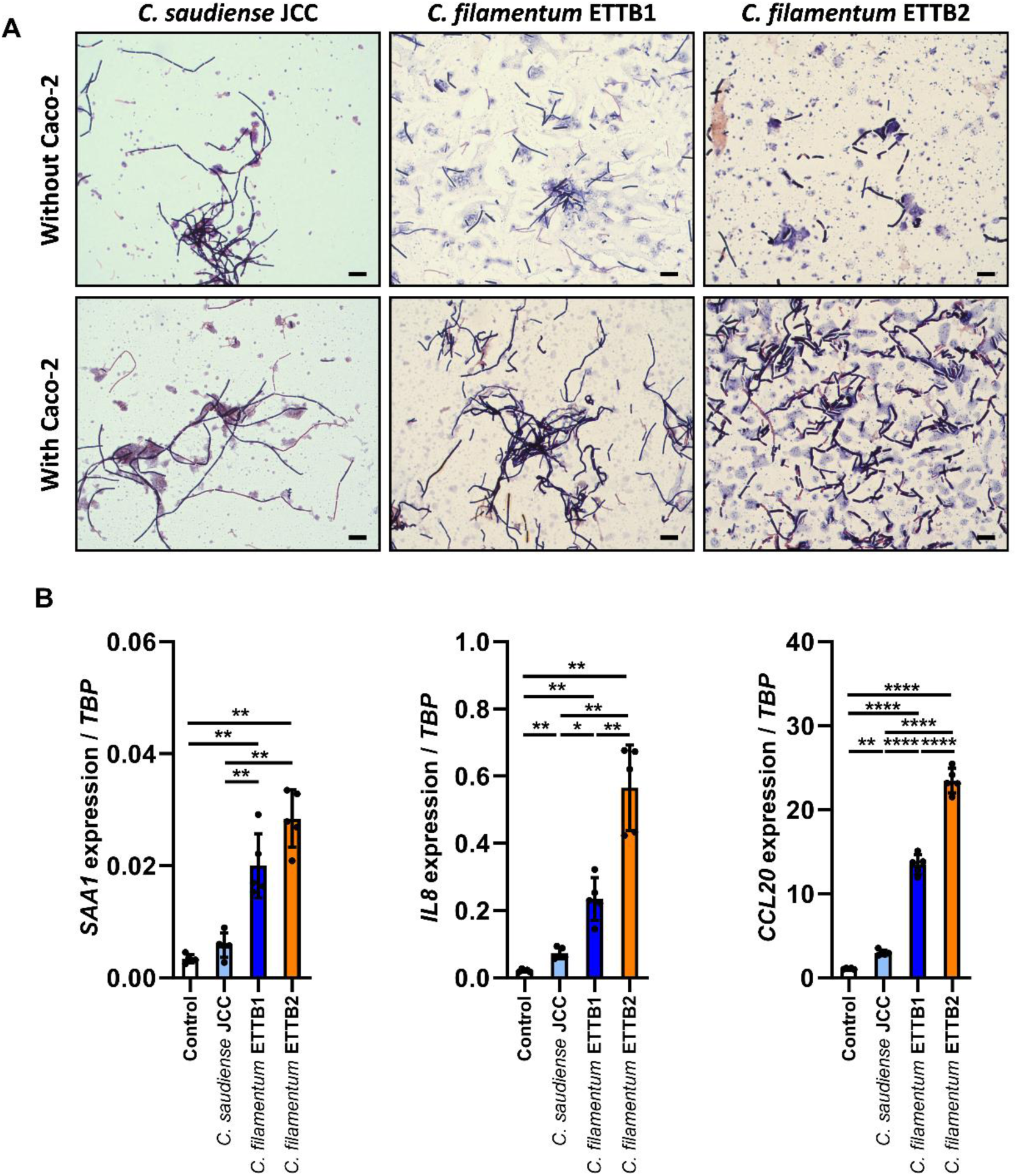
*C. filamentum* ETTB improves its growth and changes its morphology when co-cultured with Caco-2 cells, inducing epithelial innate immune responses. **A.** Representative images of *C. saudiense* JCC and *C. filamentum* ETTB1 and ETTB2 incubated in presence or absence of Caco-2 cells for 6 hours in anaerobic conditions in a Coy chamber at 37°C. **B.** Gene expression in Caco-2 cells co-cultured for 6 hours with *C. saudiense* JCC and *C. filamentum* ETTB1 and ETTB2. Scale bars represent 5 µm length. Data are presented as mean ± S.E.M. n= 5 independent experiments. One-way ANOVA with post-hoc Tukey’s test: *\*p< 0.05*, *\*\*p< 0.01* and *\*\*\*\*p< 0.0001*.

### *Clostridium filamentum* ETTB isolates express a flagellin and modulate TLR5 in Caco-2

All isolates of *C. filamentum* ETTB contained a copy of the *fliC* gene sequence in their genome (Supplementary information 2), which might encode for flagellin and interact with TLR5^12^. To investigate whether *C. filamentum* ETTB can stimulate TLR5 signaling in epithelial cells, we analysed the gene expression of *SAA1*, *IL8* and *CCL20* in the Caco-2 cells from the experiments described in Figure 4A. *C. filamentum* ETTB but not *C. saudiense* JCC increased *SAA1*, *IL8* and *CCL20* gene expression significantly compared to non-treated cells (Figure 4B). *L. reuteri* did not induce any changes when compared to control cells (Supplementary figure 3C-E), while *E. coli* did not induce *SAA1* expression but increased the expression of *IL8* and *CCL20* when compared to control non-treated cells (Supplementary figure 3F).

The gene expression of *SAA1*, *IL8* and *CCL20* has been shown to be regulated by TLR5^13,14^ and thus we next investigated if *C. filamentum* ETTB produced a protein component that activated Caco-2 cells in a TLR5-dependent fashion. First, we stimulated the cells with conditioned media (CM) from *C. filamentum* ETTB digested or not with proteinase K (CM-PK) and found that CM, but not CM-PK, of *C. filamentum* ETTB increased the gene expression of *SAA1*, *IL8* and *CCL20* (Figure 5A). We then cloned the *fliC* gene of *C. filamentum* ETTB and produced recombinant flagellin (rfliC) protein. Caco-2 cells treated with the purified flagellin protein induced *SAA1, IL8* and *CCL20* gene expression in a dose dependent manner when compared to non-treated cells (Figure 5B and Supplementary figure 4A). The gene expression induced by rfliC of *C. filamentum* ETTB was similar to cells stimulated with flagellin from *Salmonella typhimurium* (*S. typh.*), and was dependent on a functional TLR5 (Figure 5B). Consistent with these results we demonstrated that metabolically active *C. filamentum* ETTB failed to induce similar response when TLR5 was inhibited in Caco-2 cells using an antibody directed to TLR5 (iTLR5; Figure 5C).

**Figure 5:**
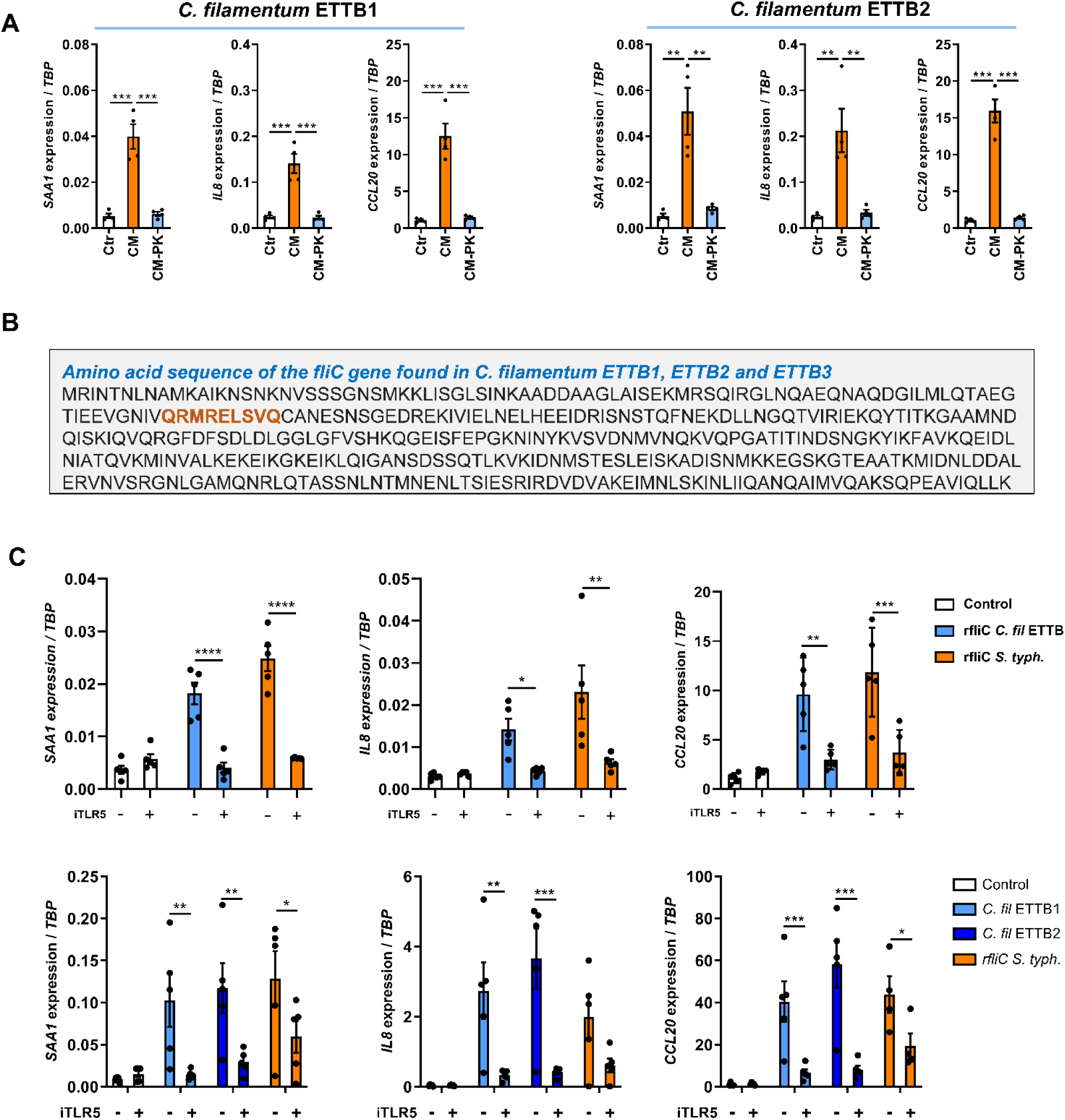
*C. filamentum* modulates the expression of innate immune genes in Caco-2 cells through fliC and the TLR5 receptor. **A.** Gene expression in Caco-2 cells incubated for 6 hours with *C. filamentum* ETTB1 and ETTB2 conditioned media (CM) or proteinase K-treated conditioned media (CM-PK). **B.** Amino acid sequence of the fliC protein encoded by the *fliC* gene of *C. filamentum* ETTB3. The TLR5 binding motif is highlighted in orange. **C.** Upper panel: gene expression in Caco-2 cells with or without TLR5 inhibition by an anti-TLR5 antibody (iTLR5) in the presence of 1 µg of recombinant fliC from *C. filamentum* ETTB3 or *Salmonella typhimurium* (*S. typh*.). Lowe panel: gene expression in Caco-2 cells with or without TLR5 inhibition by an anti-TLR5 antibody (iTLR5) after incubation for 6 hours with live *C. filamentum* ETTB1 and ETTB2 or 1 µg of recombinant fliC from *Salmonella typhimurium* (*S. typh*.). Data are presented as mean ± S.E.M. n= 4-5 independent experiments. Two-way ANOVA with post-hoc Šidák test for panel **A**. Pairwise t-test for panel **C**. *\*p< 0.05*, *\*\*p< 0.01*, *\*\*\*p< 0.001* and *\*\*\*\*p< 0.0001*.

### Prevalence and abundance of *Clostridium filamentum* ETTB3 in human cohorts

To characterize distribution of *C. filamentum* ETTB in the human gut, we screened 727 fecal metagenomes to determine prevalence and abundance of this novel species. In a longitudinal cohort of Swedish infants and their mothers, *C. filamentum* ETTB3 was highly prevalent in mothers (90% prevalence) with a median relative abundance (RA) of 0.00005 (Figure 6A), while in infants the prevalence (88%) and abundance were increasing at 12 months (median RA: 0.000365) compared to 4 months (prevalence: 15% and median RA: 0) (Figure 6A). These results indicate that *C. filamentum* ETTB is a late colonizer of the infant gut, potentially reaching higher abundances in childhood compare to adult age, as suggested by the higher abundance of *C. filamentum* ETTB3 in 12-month-old infants compared to their mothers (median RA: 0.000365 *vs.* 0.000050, *q= 0.03*) (Figure 6A).

**Figure 6:**
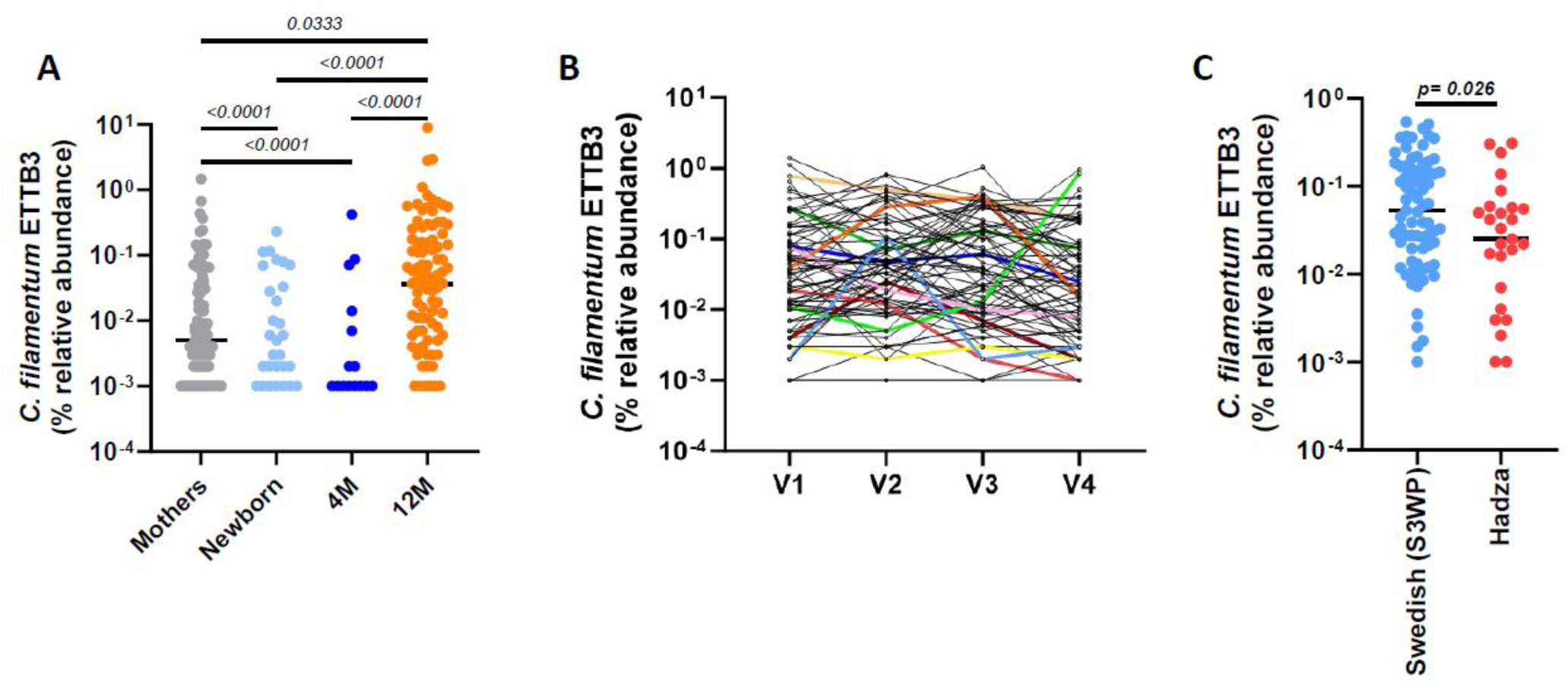
*C. filamentum* ETTB3 relative abundance in infant and adult gut microbiota. **A.** Relative abundance of *C. filamentum* ETTB3 in mothers (n=100) and infants at 0, 4 months and 12 months of age (n=100)^52^. **B.** Relative abundance of *C. filamentum* ETTB3 in a Swedish cohort of healthy individuals (S3PW) sampled 4 times over one year (n=75 individuals; 300 samples). V1, V2, V3 and V4 represent the four time points when fecal samples were assessed^33^. **C.** Relative abundance of *C. filamentum* ETTB3 in the S3WP cohort (each dot represents the average of V1-V4 relative abundances in each individual) and in Hadza hunter-gatherers of Tanzania (n= 27)^32^. Kruskal-Wallis with post-hoc Dunn’s test for panel **A** and non-parametric Mann-Whitney test for panel **C**. Adjusted P values are presented.

Next, we aimed to characterize the prevalence, abundance, and dynamics of *C. filamentum* ETTB3 in the normal adult human gut microbiota. We used metagenomics data of a Swedish cohort (S3WP), which was analysed at four time points within one year. *C. filamentum* ETTB3 was present in 100% of the individuals at all-time points. However, its abundance varied considerably between and within individuals, with an estimated RA of 0.000317 (Figure 6B and Supplementary Table 1). The abundance of *C. filamentum* ETTB3 varied over time within the same individual, with an intraclass correlation coefficient (ICC) of 0.57 (Supplementary Table 1). The variability in abundance was independent of gender (Supplementary Figure 5). We also investigated whether *C. filamentum* ETTB3 was present in a non-European cohort with a different lifestyle. In the non-industrialized hunter-gatherer population of the Hadza in Tanzania, *C. filamentum* ETTB3 had a prevalence of 96.29% and a median RA of 0.00025 (Figure 6C). The relative abundance in this population was similar to that observed in the Swedish cohorts of both adults and infants (Figure 6C).

## Discussion

In this study, we isolated and characterized three isolates belonging to a novel bacterial species, *C. filamentum* ETTB, obtained from a fecal sample of a healthy 36-year-old man. These bacteria are anaerobic and have filamentous shaped-cells. The three isolates of *C. filamentum* ETTB clustered close to a previously sequenced bacterium, *C. saudiense* JCC, isolated from a 24-year old man with morbid obesity^15^. Interestingly, genome analysis revealed that *C. filamentum* ETTB isolates have a considerably smaller genome compared to *C. saudiense* JCC and the difference in genome size may account for the metabolic differences and growth patterns observed for *C. filamentum* ETTB isolates compared to *C. saudiense* JCC, and suggests that *C. filamentum* ETTB might be a bacterium adapted to the gut.

Reduction in bacterial genome size has been linked to bacterial specialization and adaptation to specific environments or hosts^16–18^. Such processes have been observed in various bacteria, such as the *Lactobacillus* genus, *Snodgrassella alvi*, or *Candidatus Arthromitus*, suggesting mutualistic interactions between bacteria and their hosts^16,19–21^. *C. filamentum* ETTB exhibited poor growth on nutrient-rich media *in vitro*, with genome sequencing results indicating a notable reduction in the number of genes involved in carbohydrate transport and metabolism, and in amino and organic acid synthesis or degradation. More specifically, the pangenome analyses highlighted the partial or complete absence of specific pathways such as ornithine, tryptophan, vitamin B9 and B7 synthesis. There was also non-completion of modules involved in the degradation of amino acids (threonine), sugars (D-xylose and L-rhamnose), and polysaccharides (arabinan, beta-mannan, starch). Interestingly, *C. filamentum* ETTB isolates possessed complete gene sets for mannose and lactose degradation, which suggest a potential trophic specialization of *C. filamentum* ETTB in targeting specific sugars, which may positively influence host physiology. Notably, mannose supplementation in high-fat diet-fed mice has been shown to prevent obesity and metabolic disturbances, potentially by altering gut microbiota composition and function^22^. However, it remains to be explored whether *C. filamentum* prevalence or abundance in the gut affects metabolism in humans. Additionally, the pangenome analysis revealed that *C. filamentum* ETTB lacked the module to synthesize D-lactate, which is in concordance with our *in vitro* measurements of organic acid production showing that *C. filamentum* ETTB produced less lactate compared to *C. saudiense* JCC. However, *C. filamentum* ETTB produced similar amounts of acetate, suggesting that in *C. filamentum* ETTB the main product of fermentation is acetate and not lactate.

To further illustrate the potential specialization of *C. filamentum* ETTB, we observed a significant increase in the number of genes associated with cell motility, particularly the presence of the *fliC* gene, encoding a flagellin protein. Flagella are crucial cell motility structures enabling bacteria to migrate or interact with epithelial cells in a host-specific manner^23,24^. This interaction between host cells and bacteria can trigger immune or inflammatory responses through the binding of flagellin and more specifically fliC protein to host receptors such as TLR5^12,25^. Our results showed that *C. filamentum* ETTB increased the expression of *SAA1*, *IL8*, and *CCL20* in Caco-2 cells, which required TLR5 signaling and was mediated by the fliC protein. Additionally, exposure of *C. filamentum* ETTB to Caco-2 cells resulted in increased bacterial biomass and filamentation. Interestingly, *C. saudiense* JCC did not increase in biomass or become more filamentous when exposed to Caco-2 cells and its genome lacked a copy of a *fliC* gene. This might explain the absence of induction of *SAA1*, *IL8* and *CCL20* expression in Caco-2 cells when grown in the presence of *C. saudiense* JCC. Several studies have suggested that bacterial filamentation reflects an adaptive response to specific environments, aiding in surface colonization, growth, and metabolic optimization^26–28^. These findings parallel studies investigating the function of segmented filamentous bacteria, particularly *Candidatus Arthromitus*, highlighting a unique interaction with host cells^26^.

*Candidatus Arthromitus*, possessing a small genome size of around 1.6Mb, relies on other bacteria and host factors for growth^29,30^. It is also recognized as a key gut bacterium capable of inducing immune system maturation in rodents, particularly Th17 lymphocyte development, partly through *Saa1* and *Saa3* expression^23,26,31^. Although our results suggest potential similarities for the ability of *C. filamentum* ETTB to change its morphology in contact with host cells and to interact with host immune responses, it remains to be determined whether *C. filamentum* ETTB isolates rely on other bacterial members of the human gut or host cells to enhance their growth, induce TLR5 activation and promote immune system maturation in the gut *in vivo*.

We also analysed the prevalence, abundance, and dynamics of *C. filamentum* ETTB in three human cohorts and found that this bacterium is highly prevalent in the gut of both industrialized and non-industrialized populations. The abundance of *C. filamentum* ETTB was similar in the Swedish S3WP cohort and the Hadza hunter-gatherers, suggesting that this bacterium is prevalent in healthy humans and may be associated with non-westernized or fiber-rich diets^32^.

Additionally, *C. filamentum* ETTB can be considered a late colonizer, exhibiting higher relative abundance at 12 months after birth. This suggests that *C. filamentum* ETTB may require specific gut conditions, such as an anoxic environment and a complex microbiota potentially, as its genetic makeup indicates metabolic dependency.

Our results also demonstrated that the abundance of *C. filamentum* ETTB is variable over a one-year period in healthy Swedes from the S3WP, with significant inter- and intra-individual variability. While the factors driving variation in the fecal levels of *C. filamentum* ETTB remain unclear, we speculate that the abundance and fluctuations of this bacterium may be influenced by the availability of nutrients or dietary intake, as well as the dynamics of other bacteria that may cross-feed with *C. filamentum* ETTB^33^.

In summary, our findings suggest that *C. filamentum* ETTB may be a specialized gut bacterium, potentially requiring specific interactions with host cells or microbial niches for optimized growth.

## Materials and Methods

### Isolation of *Clostridium filamentum* ETTB isolates from human feces

A simulated human intestinal redox model (SHRIM) was used to isolate ETTB strains from a stabilized gut microbial community under *in vitro* conditions. SHRIM is a two-chamber bioreactor with an anaerobic luminal compartment (250 mL) and an oxygen feeder (100 mL), which are separated by an agar-mucin coated Nafion Membrane N115 (DuPont, USA; diameter 2.5 cm) and continuously purged with nitrogen and oxygen, respectively (Supplementary Figure 1A). This condition mimics the physiological oxygen diffusion gradient at the gut mucosa. The agar-mucin coated Nafion Membrane was prepared by dissolving 2% agar, 0.5% mucin type II in phosphate buffer saline and was autoclaved. After the autoclave pH was adjusted to ∼7 and molten agar mucin mixture overlayed (1cm) on the Nafion septum facing the anaerobic luminal compartment. The agar-mucin was allowed to solidify before connecting the two compartments of the SHRIM bioreactor. The luminal chamber was continuously stirred at 250 rpm. and kept at 37 °C. The oxygen feeder contained 100mM potassium phosphate buffer. The luminal chamber was seeded with 250 mL of feed containing: (in g/L) arabinogalactan (1.0), pectin (2.0), xylose (1.5), starch (3.0), glucose (0.4), yeast extract (3.0), peptone (1.0), mucin type II (4.0), and cysteine (0.5). To simulate digestion processes, the feed was acidified to around pH 2 with 6M HCl and neutralized with simulated pancreatic juice to a pH of around 6.9. The simulated pancreatic juice contained: (in g/L) NaHCO3 (12.5), Oxgall bile salts (6.0), and pancreatin (0.9). The feed and pancreatic juice mix (70:30), referred to as SHRIM feed, was kept anaerobic by continuously purging with nitrogen^34^. The SHRIM feed was fed continuously to the luminal chamber at a rate giving a retention time of around 24 h, and pH was maintained between 6.6 and 6.9 with pH controller and dosing Pump (Black stone BL7912, Hanna Instruments, UK).

The SHRIM system was inoculated with an aliquot of the freshly voided fecal sample from a healthy donor. A preculture was prepared anaerobically in a Coy chamber (5% hydrogen, 10% carbon dioxide, and 85% nitrogen) by adding 2% (w/v) fecal material to 5 ml of LYBHI broth. The LYBHI broth was prepared by dissolving (g/L): BHI (37), yeast extract (5), cellobiose (1), maltose (1), cysteine (0.5) and 3ml/L of hemin solution (0.2%). The preculture was incubated for 3 h at 37 °C, and 2% of the pre-culture was seeded into the luminal compartment of the SHRIM. The SHRIM bioreactor was run in continuous mode for 1 week. After this period, the SHRIM bioreactor was moved to the Coy chamber. The luminal compartment was emptied, and the agar-mucous septum was removed for Gram staining and subculturing to isolate sessile bacterial strains. The luminal-facing agar-mucin surface was scraped with a loop and inoculated into LYBHI broth. Based on peculiar morphology, ETTB strains were isolated through repeated subculturing and pure culture techniques using LYBHI agar plates. After confirming the purity, pure cultures of ETTB strains were preserved in LYBHI medium containing 20% glycerol at -80oC.

### Bacterial cultures

*C. saudiense* JCC (DSM 27835), *C. filamentum* ETTB1 (DSM 117630), ETTB2 and ETTB3 (DMS 115107, CCUG 77667), *Lactobacillus reuteri* and *Escherichia coli* were cultured anaerobically in autoclaved LYBHI, containing brain-heart infusion medium (BHI from Oxoid, ref. CM1135) supplemented with yeast extract (0.5%, Oxoid, ref. LP0021), hemin (0.005%, Sigma-Aldrich, ref. 51280-5G), L-cysteine (0.05%, Sigma-Aldrich, ref. W326305-100G), maltose (0.1%, Millipore, ref. 1.05910.0500) and D-(+)-cellobiose (0.1%, Sigma-Aldrich, ref. 22150-50G). All bacterial incubations were performed in strict anaerobic conditions with 10% CO2 and 5% hydrogen inside a Coy chamber at 37°C.

### Gram staining

Bacterial isolates were regularly checked for purity using Gram-staining. Cells were dried on a clean slide and stained with crystal violet (0.5%) and/or iodine (0.5%) and safranin (0.5%) reagents. Pictures were acquired using a light microscope with a magnification of 100x (Axio Lab.A1).

### Scanning electron microscopy of *Clostridium saudiense* JCC and *Clostridium filamentum* ETTB isolates

Bacterial suspensions were allowed to sediment on silica wafers for 30 minutes followed with a fixation using Karnovsky fixative in cacodylate buffer for one hour at room temperature. The samples were treated with osmium (1%) for 30 minutes at room temperature followed with several steps of washing using water. The samples are dehydrated using serial dilutions of ethanol (30%, 1x50, 70, 85, 95, 100%) baths with 5 minutes incubation each and treated with hexamethyldisilazane for 2 minutes. The samples were coated with gold with total thickness of 2 nm. Images were acquired using the scanning electron microscope (Zeiss Gemini 450) with electron high tension of 2.00 KV and magnifications of 7 and 25 KX.

### Metabolic phenotyping of *Clostridium saudiense* JCC and *Clostridium filamentum* ETTB isolates

Metabolic phenotyping of the isolates was performed using analytical profile index assays (API® ZYM (ref. 25200), API® 20A (ref. 20300) and rapid ID 32 STREP (ref. 32600), Biomerieux). The analyses were performed on *C. saudiense* JCC, *C. filamentum* ETTB1, ETTB2 and ETTB3. Fresh cultures from 2-3 LYBHI agar plates (Bacto Agar, ref. 214010, BD) were used for the analysis. A bacterial suspension was prepared according to manufacturer’s instructions in a suspension media (API sodium chloride (NaCl 0.9%) or API 20A medium). The density of 3 to 5 following the McFarland standards was used. The tests were performed using three independent replicates in a Coy anaerobic chamber at 37°C. The results were obtained after an incubation of 24 hours for API® 20A and 4 hours for API® ZYM and rapid ID 32 STREP kits.

### Growth assessments of Clostridium saudiense JCC and Clostridium filamentum ETTB isolates

For each isolate, a single colony from LYBHI plates (2% agar) incubated anaerobically in a Coy chamber for 24h was collected using a sterile loop and inoculated in 3 mL of LYBHI media in 15 mL tubes. Bacterial suspensions were sampled at different time points: 0, 2, 4, 6, 8, 12, 24 and 48 hours. Bacterial growth was estimated by quantifying using a spectrophotometer at optical density of 600 nm.

### Short-chain fatty acids analysis

Short-chain fatty acids were measured using gas chromatography coupled to mass spectrometry detection (GC-MS) as described previously^35^. Briefly, approximately 25 µL of overnight culture media were mixed with internal standards, and added to glass vials. All samples were then acidified with HCl, and SCFAs were extracted with two rounds of diethyl ether extraction. The organic supernatant was collected; the derivatization agent N-tert-butyldimethylsilyl-N-methyltrifluoroacetamide (Sigma-Aldrich) was added and samples were incubated at room temperature overnight. SCFAs were quantified with a gas chromatograph (Agilent Technologies 7890A) coupled to a mass spectrometer (Agilent Technologies 5975C). Short-chain fatty acid standards were attained from Sigma-Aldrich (Stockholm, Sweden).

### DNA extraction and analysis of the 16S rRNA gene

Bacteria were cultured overnight in LYBHI media. 3ml aliquots of each culture were centrifuged, washed, and suspended in lysis buffer SL2 from the Nucleospin® Soil kit (Macherey-Nagel, ref. 740779). Cells were sheared with 3 rounds of bead beating for 1 minute at 5.0 m/s in a FastPrep®-24 Instrument (MP Biomedicals) and DNA was extracted using the Nucleospin® Soil kit. DNA concentrations were estimated by Nanodrop. Amplification of bacterial 16S rRNA genes was performed using the 27F and 1492R universal primers. PCR products were purified using NucleoSpin® Gel and PCR Clean-up (Macherey-Nagel, ref. 740609.250) and the concentrations were estimated using Quanti-iT^TM^ PicoGreen® dsDNA Assay Kit (Invitrogen, ref. P11496). The sequencing of the PCR products was performed using Eurofins services.

### DNA extraction for whole genome sequencing

High-quality DNA was extracted using a modified version of the Marmur procedure^36^. Overnight bacterial cultures were centrifuged and resuspended in 800 µL of sterile EDTA-saline solution and vortexed at maximum speed to mix thoroughly. Cells were lysed using 10 µL of lysozyme (300 mg/mL), 10 µL of mutanolysin (1000 U/mL) and 10 µL of RNase A (100 mg/mL) at 37°C for 45 minutes and vortexed every 15 minutes. After the first incubation, 80 µL of sodium dodecyl sulfate (SDS) 25% was added and the samples were vortexed and incubated at 65 degrees for 10 minutes. A volume of 250 µL of sodium chloride (5 M) was added to the samples and vortexed at maximum speed for 10 seconds followed by an addition of 400 µL of chloroform:isoamyl alcohol and shaking for 15 minutes at 1400 rpm using an orbital shaker. The samples were centrifuged for 15 minutes at 4 degrees at 14000 rpm and the aqueous phase was transferred to a new tube. For each 1 mL of aqueous phase 90 µL of sodium acetate (3 M) was added followed by 600 µL of cold isopropanol. The samples were centrifuged for 10 minutes at 13000 g and the supernatant was discarded to allow the DNA to dry completely. The DNA pellet was dissolved in Low TE buffer and incubated at 4 degrees overnight for complete resuspension.

### Genome sequencing

DNA integrity and concentration were evaluated using Tapestation 4150 with Genomic DNA ScreenTape and reagents (Agilent) and Qubit 3.0 Fluorometer and Qubit dsDNA BR assay kit (ThermoFisher Scientific). For Illumina short-reads sequencing, libraries were prepared using the TruSeq DNA PCR-free Library Preparation kit (Illumina, 20015963 and 20015949) after fragmentation with a Covaris S220 Focused-ultrasonicator (Covaris) generating 550-bp inserts. Libraries were quantified using Quant-iT dsDNA HS Assay Kit (ThermoFisher Scientific) and sequenced on an Illumina Miseq instrument using MiSeq Reagent Kit v3, 600 cycles. For Pacbio sequencing, genomic DNA was assessed using the pulse field electrophoresis (Pippin), DNA was sheared using g-TUBE (Covaris, 10145) and cleaned with AMPure XP beads (Beckman Coulter, A63882); libraries were prepared using the SMARTBell template Prep kit 1.0SPv3 (101-991-900) with the Barcoded adaptor kit 8A (101-081-300) and quality controlled using a Fragment Analyzer (Agilent) before being sequenced on a Sequel sequencer for 10 hours. After demultiplexing, samples had a mean 663,538 read number with a ∼4 kb read length. Assemblies were performed using the LORA pipeline v1.0.0 from Sequana^37^. From the demultiplexed samples (mean read count per sample: 663,538), the LORA pipeline was run with default parameters and reproducible containers from the Damona project (https://github.com/cokelaer/damona). In particular, the assembly was performed with Canu^38^. Circularization was performed as (Hunt et al., 2015)^39^, and coverage homogeneity was assessed using the Sequana Coverage software for quality control^40^. The final scaffolds each consisted of a single circularized contig of 2.8 Mb with a mean fold read coverage of 72 to 124 per sample.

For Oxford Nanopore long-reads sequencing, genomic DNA was prepared using the Rapid barcoding kit (SQK-RBK004) following the manufacturer’s instructions (ONT) and sequenced on a ONT’s MinION device on a R9.4.1 flow cell (FLO-MIN106D). Base-calling was performed using ONT’s guppy v. 4.2.2. Genomes were obtained by hybrid assembly of Nanopore and Illumina reads. The Unicycler pipeline v0.4.8 in hybrid mode used to obtain de novo assemblies. All dependencies for Unicycler were installed in a conda environment. The dependency programs include SPAdes v3.13.0, racon v1.4.1, bowtie2 v2.3.5.1, and pilon v1.23. The hybrid assemblies were annotated using Prokka v1.14.5 (https://github.com/tseemann/prokka).

### Average nucleotide identity and phylogenetic tree analysis

Sequences in Fasta format have been uploaded at the Bacteria and viral Bioinformatics Resource Center (BV-BRC, https://www.bv-brc.org/) and analysed using the phylogenetic tree tool. Basically, nucleotides are aligned throughout mafft alignment using 27 genomes and 78 single-copy genes and proteins^41^. The analysis produced a RAxML using an LG model. ANI values have been estimated using an online ANI calculator tool that adopts the OrthoANIu algorithm developed by Yoon et al. (2017)^42^.

### *In silico* validation and description of novel taxa

The sequences of 16S rRNA and the whole genome of *C. filamentum* ETTB1, ETTB2, ETTB3 and *C. saudiense* JCC were analysed using Protologger tool^11^, Galaxy version 1.0.0 (https://protologger.bi.denbi.de/).

### Pangenome analysis

Genome annotation was performed using DRAM^43^, an ensemble annotation tool that integrates multiple databases, including PFAM^44^, KEGG^45^, CAZY^46^, and MEROPS^47^. The initial genome annotations were distilled through distillR^48^, an R package that leverages KEGG^45^ and MetaCyc^49^ metabolic pathways to compute genome-inferred functional traits, providing insights into each genome’s potential for compound degradation or synthesis. These refined annotations were used to evaluate the metabolic capabilities of all genomes under analysis and to detect gene-level variations between different strains.

Next, a pangenome graph encompassing three Clostridium filamentum genomes and C. saudiense was constructed using pggb^50^. The pangenome graph clusters identical sequences shared among genomes, while genome-specific regions are represented as distinct branches. After constructing the graph, pangenome visualizations were generated using odgi^51^.

### Co-culture of Caco-2 cells with *Clostridium saudiense* JCC and *Clostridium filamentum ETTB* isolates

Caco-2 clone HTB37 cells (passage between 17 and 20) were used for co-culture experiments. The cells were checked regularly for mycoplasma contaminations and maintained in IMDM Hyclone medium supplemented with 10% foetal bovine serum and 1% Penicillin/streptomycin. Co-culture experiments were performed in 12-well plates in anaerobic conditions in a Coy chamber with incubation at 37°C. Caco-2 cells were seeded at 1x10^5^ cells per well and used for co-culture once a monolayer was formed. The media used during the co-culture was composed of 80% IMDM without antibiotics and 20% LYBHI.

To block TLR5 activity, Caco-2 cells were incubated for one hour with anti-hTLR5 (maba2-htlr5, InvivoGen) antibody before starting the co-culture with bacterial isolates. After one hour of incubation, Caco-2 cells were washed twice with phosphate buffered saline before starting the incubation with bacterial isolates.

### Quantification of colony forming units

Bacteria were collected from the co-culture experiments after 6 hours of incubation and centrifuged in sterile tubes. The supernatants were discarded in anaerobic conditions in a Coy chamber, and the bacterial pellets were resuspended in 50 µL of fresh LYBHI media. Cellular suspensions were serially diluted in LYBHI media and plated in duplicate on LYBHI agar plate. Colonies were counted 24 hours after incubation at 37°C under anaerobic conditions in a Coy chamber.

### Preparation of bacterial conditioned media

To produce bacterial conditioned media, a single colony for each isolate was collected from a 24-hour culture plate (LYBHI with 2% agar) and cultured in 3 mL of LYBHI media and incubated overnight at 37°C. The bacterial cultures were centrifuged at 4°C for 10 minutes at 4500 rpm and the supernatants were collected and filtered using sterile 0.22 µm filters. Part of the conditioned-media was digested using proteinase K (50 microg/mL) for 30 minutes at 37°C followed by 5 minutes incubation at 85°C for enzymatic activity inactivation. Additionally, the bacterial isolates were heat-killed at 65°C for 30 minutes. Bacterial viability after the heat-killing was assessed using agar plates incubated in anaerobic conditions at 37°C, showing no viable colonies after heat-killing. Freshly prepared conditioned media was used in all co-culture experiments with Caco-2 cells.

### Preparation of recombinant fliC protein of *Clostridium filamentum*

The *fliC* gene sequence of *C. filamentum* ETTB was cloned into the pETM11 vector, which carries a kanamycin resistance cassette. For protein expression, ClearColi cells (Lucigen) were transformed with 1 µl plasmid DNA. Cells were cultured overnight at 37°C in LB medium supplemented with 50 µg/mL kanamycin, then diluted 1:50 into 400 mL fresh media the following day. When optical density (OD600) reached 0.4-0.6/mL, protein expression was induced with the addition of 0.1 mM IPTG. Following incubation at 37°C for two hours, cells were harvested by centrifugation at 5600 rpm for 15 minutes. The pellet was resuspended in 8 mL urea lysis buffer (100 mM NaH2PO4, 10 mM Tris-HCl, 8 M urea, pH 8) at room temperature for 2 hours. Lysate was cleared by centrifugation at 25,000 x g for 20 minutes and supernatant was incubated with 4 mL Ni-NTA agarose at room temperature for 2 hours to facilitate protein binding. FliC-bound Ni-NTA slurry was washed once with 5 mL urea lysis buffer pH 8, then twice with urea lysis buffers of decreasing pH (pH 6.3 and pH 5.9). Protein was eluted with urea lysis buffer pH 4.5 in 500 µL fractions and analyzed by SDS-PAGE. FliC-containing fractions, as determined by Coomassie stain, were pooled and dialyzed against 20 mM Tris-HCl (pH 8), 300 mM NaCl, and 5 mM MgCl₂ using 10 kDa molecular weight cut-off dialysis cassettes (Pierce). Protein concentration was determined by BCA (Pierce) prior to storage at -80C.

### RNA extraction from Caco-2 cells and tissues

Caco-2 cells were washed with cold PBS and lysed with TriPure reagents (ref. Sigma-Aldrich, 11667167001). RNA extraction was performed using the Trizol/Chloroform:isoamyl method and the RNeasy Kit (Qiagen, ref. 74106).

### Gene expression analysis

cDNA synthesis was performed on 400 ng of RNA for Caco-2 cells, using High-Capacity cDNA Reverse Transcriptase kit (Applied biosystems, ref. 4368813). qPCR analyses were performed using the iQ SYBR Green supermix (Bio-Rad, ref. 1708886). Tata-binding Protein (*TBP*) gene was used as a housekeeping gene.

### Statistical analysis

Statistical analysis was performed using GraphPad Prism V.10. Mann-Whitney test was used for pairwise comparisons and One-way ANOVA with Tukey’s post hoc analysis was used for parametric ANOVA between groups. The Kruskal-Wallis with post-hoc Dunn’s method was used for non-parametric ANOVA between groups. To evaluate the mean differences between groups having two independent variables, Two-way analysis of variance (ANOVA) with post-hoc Šidák’s test was performed. Data are presented as mean ± SEM unless otherwise noted.

## Supporting information

Makki et al_suppdata

## Acknowledgements

We are grateful to the Centre for Cellular Imaging at the University of Gothenburg for providing assistance in microscopy and Professor Gunnar Hansson for providing the Caco-2 cell line. These studies have been funded by the Wallenberg Foundation (2017.0026), the Swedish Heart and Lung Foundation (20210366), the Novo Nordisk Foundation the Swedish Research Council (2019-01599), and grants from the Swedish state under the agreement between the Swedish government and the county councils, the ALF-agreement (ALFGBG-718101). The computations were enabled by resources in projects [SNIC 2022/5-451 and NAISS 2023/5-389] provided by the National Academic Infrastructure for Supercomputing in Sweden (NAISS) and the Swedish National Infrastructure for Computing (SNIC) at UPPMAX, funded by the Swedish Research Council through grant agreements no. 2022-06725 and no. 2018-05973. PS is funded through the Bettencourt Coup d’elan, the Bill and Melinda Gates Foundation Grand Challenge Grant OPP11413338 and ERC CoG Nicheadapt.

## Author contributions

Conceptualization: K.M., V.T., and F.B.; Investigation: K.M., V.T., M-E.M., M-T.K.; Methodology: K.M., A.Q., A.A, O.A., M-E.M., V.T., M-T.K., S.C., C.F., G.Y., P-O.B., P.S., R.L., H.B., C.D., J.J. and F.B.; Resources, R.L., P.S., F.B.; Writing – Original Draft, K.M., V.T, and F.B. Writing – Review & Editing: K.M., A.Q., M-E.M., A.A., O.A., P.S., M-T.K., S.C., R.L., C.D., V.T. and F.B.; Supervision: K.M., V.T. and F.B.

## Supplementary figures

**Supplementary figure 1:**

**A.** Cartoon representing a simulated human intestinal redox model (SHRIM) and describing the isolation procedure used to isolate *C. filamentum* ETTB1, ETTB2 and ETTB3. **B**. Gram staining of *C. saudiense* JCC and *C. filamentum* ETTB1, ETTB2 and ETTB3. Colonies were collected after 24 hours of growth on LYBHI agar plates in anaerobic conditions at 37°C. **C.** Polar lipid profiles of *C. saudiense* JCC and *C. filamentum* ETTB1, ETTB2 and ETTB3.

**Supplementary figure 2:**

**A.** Linear pangenome analyses highlighting genomic structural differences between *C. saudiense* JCC and *C. filamentum* species. **B.** Comparative analysis of Clusters of Orthologous Groups (COG) for *C. saudiense* JCC and *C. filamentum* ETTB3 highlighting genomic functional differences in the two bacterial species. Data are presented in absolute number (left panel) and percentage (right panel).

**Supplementary figure 3:**

**A.** Representative images of *C. filamentum* ETTB1 and ETTB2, *Lactobacillus reuteri* (*L. reuteri* or Lr) and *Escherichia coli* (*E. coli* or Ec) when incubated in presence or absence of Caco-2 cells for 6 hours under anaerobic conditions at 37°C. Scale bars represent 5 µm length. **B.** Colony forming unit counts (CFU/ml) of *C. filamentum* ETTB1 and ETTB2, *Lactobacillus reuteri* (*L. reuteri* or Lr) and *Escherichia coli* (*E. coli* or Ec) after 6 hours of incubation in presence or absence of Caco-2 cells under anaerobic conditions at 37°C. Gene expression in Caco-2 cells when incubated for 6 hours with **C.** *C. filamentum* ETTB1, **D.** *C. filamentum* ETTB2, **E.** *L. reuteri*, **F.** *E. coli.* Data are presented as mean ± S.E.M. n= 5-6 independent experiments. Mann-Whitney: *\*\*p< 0.01*, *\*\*\*p< 0.001*.

**Supplementary figure 4:**

**A.** Gene expression in Caco-2 cells treated with different concentrations of the recombinant fliC protein from *C. filamentum* ETTB isolates. Data are presented as mean ± S.E.M. n= 3 independent experiments. One-way ANOVA with post-hoc Dunnett’s test: *\*\*p< 0.01*, *\*\*\*p< 0.001*, *\*\*\*\*p< 0.0001*.

**Supplementary figure 5:**

Relative abundance of *C. filamentum* ETTB3 for men and women in the Swedish S3PW cohort sampled 4 times over a one-year period (n=75 individuals; 300 samples). V1, V2, V3 and V4 represent the four time points when fecal samples were assessed^33^.

