## Supplementary material for "Characterization and description of *Clostridium filamentum* ETTB, a novel gut bacterium with TLR5 modulating properties": Makki et al_suppdata

**A**

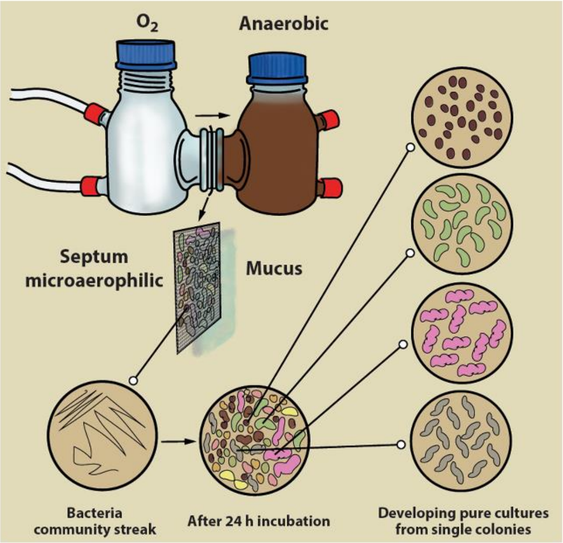


**B**

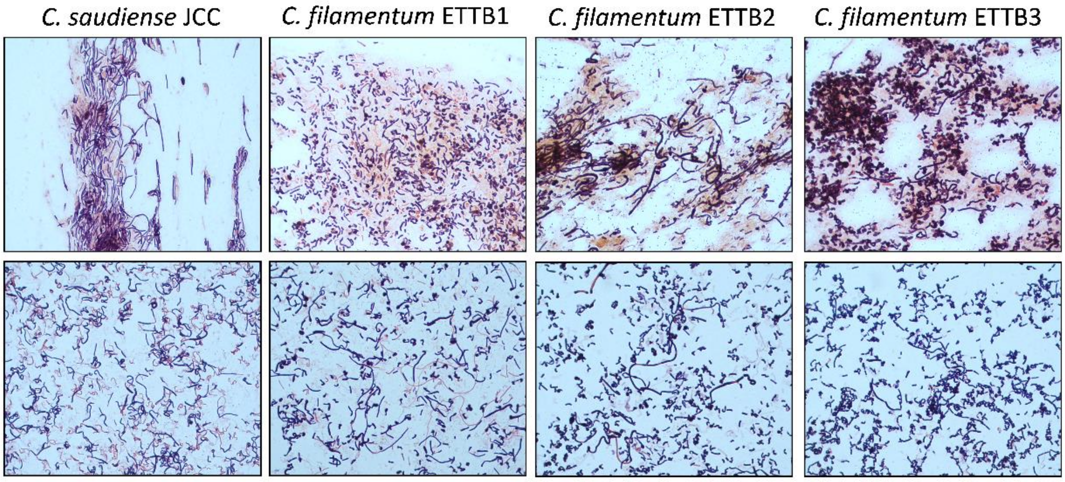


**Gram staining: bacteria collected from LYBHI agar plates after 24 hours culture under anaerobic conditions**

**C**

### Polar lipid profile

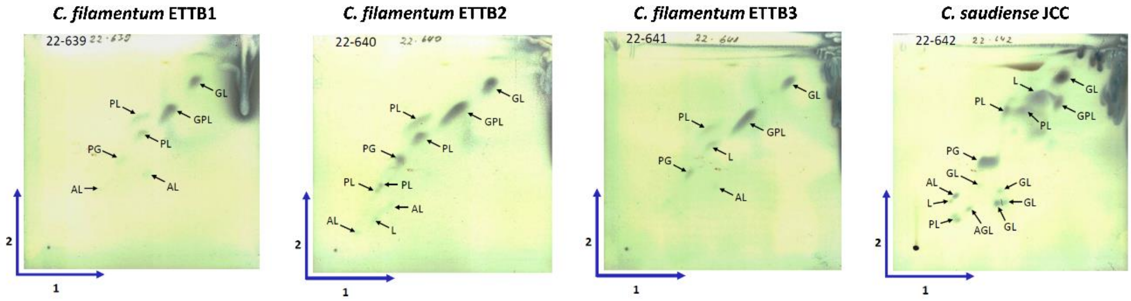


PG = Phosphatidylglycerol, GPL = Glycophospholipid, AGL = Aminoglycolipid, AL = Aminolipid, GL = Glycolipid
PL = Phospholipid, L = Lipid

**Supplementary figure 1**

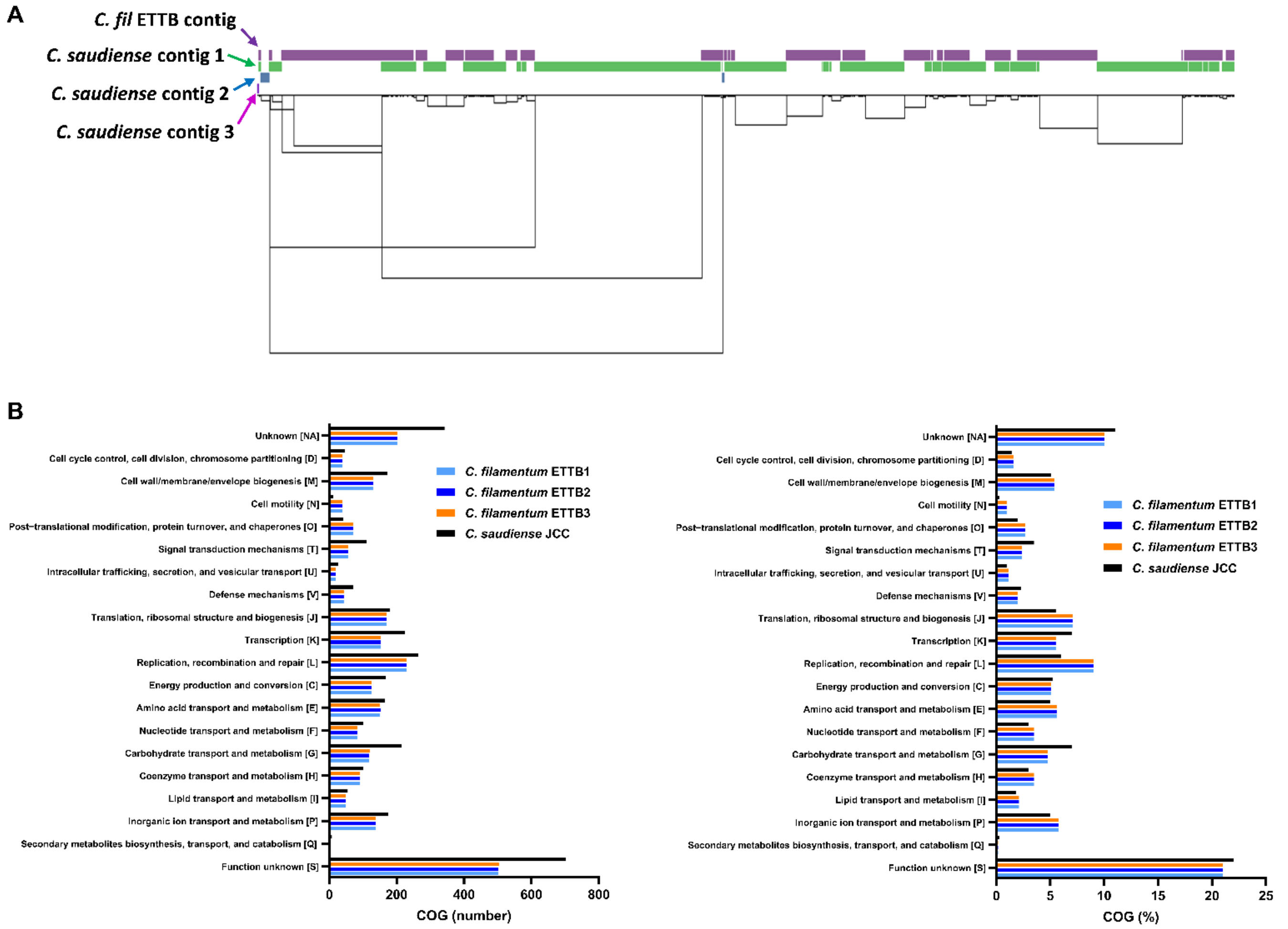


**Supplementary figure 2**

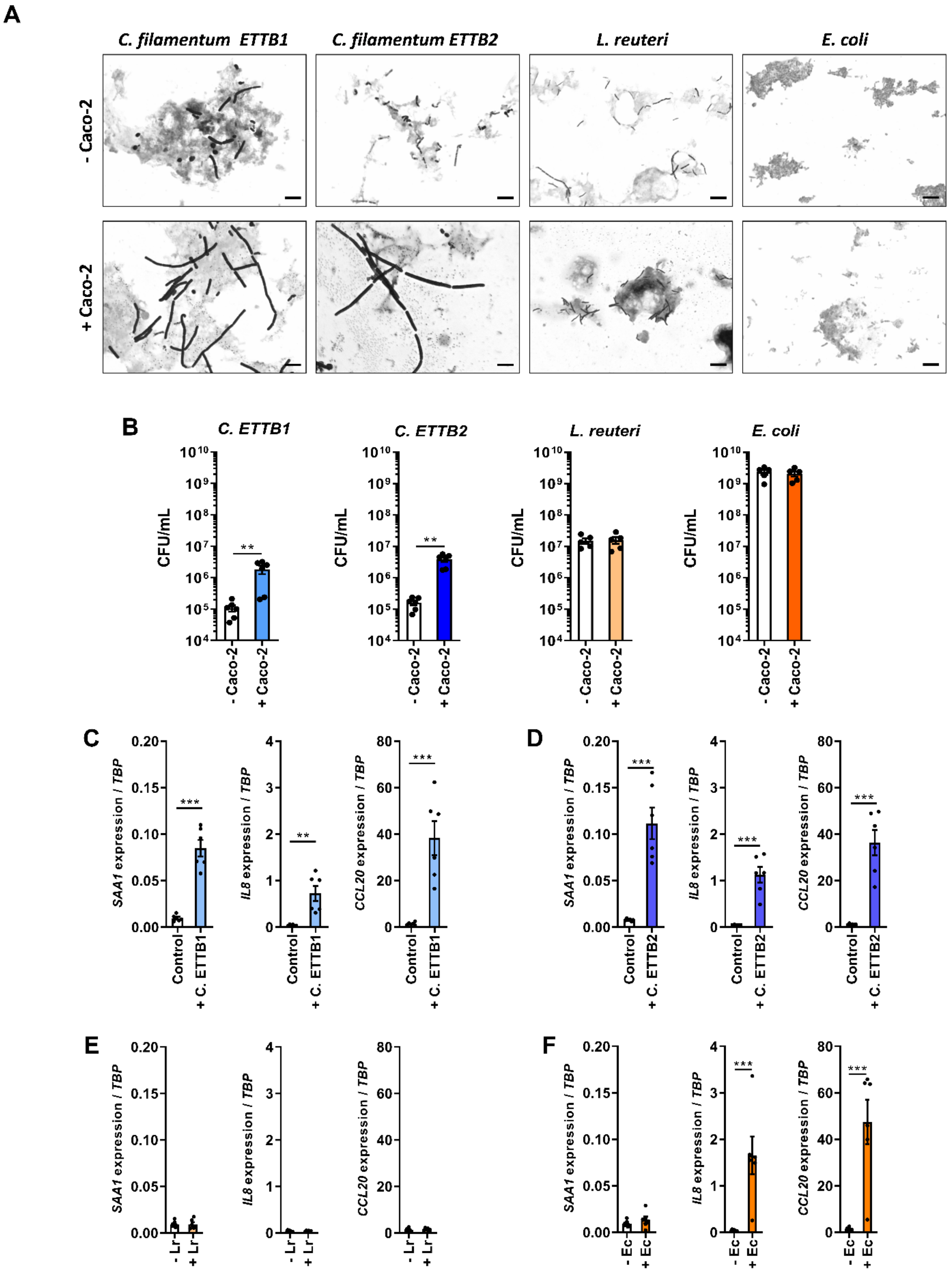


Supplementary figure 3

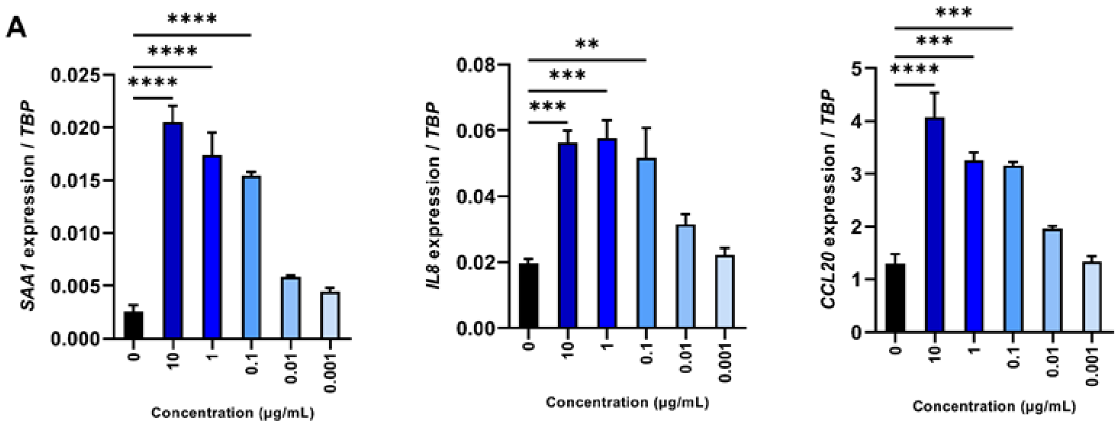


Supplementary figure 4

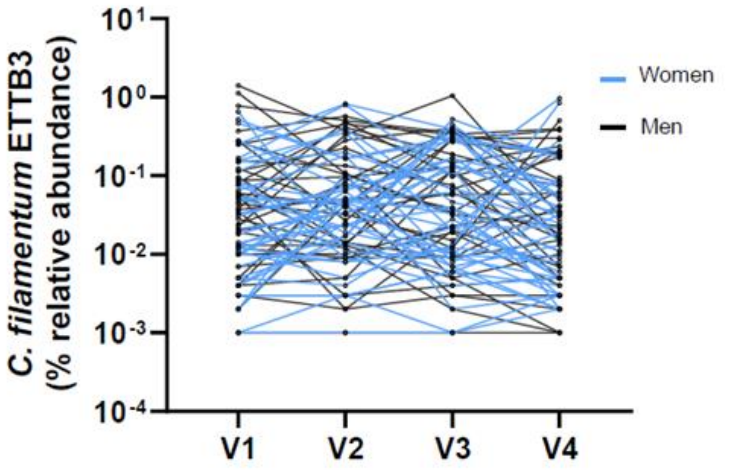


Supplementary figure 5

**Supplementary table 1**

| | *C. filamentum* ETTB3 |
|---|---|
| **log_Estimate** | -3.498499 |
| **Estimate_relab** | 0.0003173226 |
| **Variance_between** | 0.3233905 |
| **Variance_within** | 0.2454845 |
| **Variance_total** | 0.568875 |
| **ICC** | 0.5684737 |
| **Index_of_individuality_COUNTS** | 0.7590962 |
| **SD_total** | 0.754238 |
| **SD_between** | 0.5686743 |
| **SD within** | 0.4954639 |
| **Pvalue_within** | 0.623843 |
| **2,50%** | -4.958607 |
| **97,50%** | -2.146582 |
| **range** | 2.812025 |

**Supplementary information 1**

```
CLUSTAL 2.1 Multiple Sequence Alignments


Sequence type explicitly set to DNA
Sequence format is Pearson
Sequence 1: NR_144696.1 C. saudiense JCC  1478 bp
Sequence 2: ettb3        1420 bp
Sequence 3: ettb1        1416 bp
Sequence 4: ettb2        1422 bp
Start of Pairwise alignments
Aligning...

Sequences (1:2) Aligned. Score:  97
Sequences (1:3) Aligned. Score:  97
Sequences (1:4) Aligned. Score:  97
Sequences (2:3) Aligned. Score:  98
Sequences (2:4) Aligned. Score:  97
Sequences (3:4) Aligned. Score:  97
Guide tree file created:   [clustalw.dnd]

There are 3 groups
Start of Multiple Alignment

Aligning...
Group 1: Sequences:   2      Score:26306
Group 2: Sequences:   3      Score:26214
Group 3: Sequences:   4      Score:26041
Alignment Score 58624

CLUSTAL-Alignment file created  [clustalw.aln]
```

---

clustalw.aln

```
CLUSTAL 2.1 multiple sequence alignment


ettb3            -------------------------------GGAACGGGGTGCTAACATGCA-GTCGAGC
ettb1            ---------------------------------CGTACGCTGCTAACCTGCA-GTCGAGC
NR_144696.1      AGAGTTTGATCCTGGCTCAGGACGAACGCTGGCGGCGTGCCTAACACATGCAAGTCGAGC
ettb2            ------------------------------CAAGGCG-GCTGCTAAAATGCA-GTCGAGC
                                                       *      *  **** *******

ettb3            GAGTGAATCTCCTTCGGGAGTGAAGCTAGCGGCGGACGGGTGAGTAACACGTGGGCAACC
ettb1            GAGTGAATCTCCTTCGGGAGTGAAGCTAGCGGCGGACGGGTGAGTAACACGTGGGCAACC
NR_144696.1      GAGTGAATCTCCTTCGGGAGTGAAGCTAGCGGCGGACGGGTGAGTAACACGTGGGCAACC
ettb2            GAGTGAATCTCCTTCGGGAGTGAAGCTAGCGGCGGACGGGTGAGTAACACGTGGGCAACC
                 ************************************************************

ettb3            TGCCTCATAGAGGGGAATAGCCTTTCGAAAGGAAGATTAATACCGCATAAGATTGTAGCT
ettb1            TGCCTCATAGAGGGGAATAGCCTTTCGAAAGGAAGATTAATACCGCATAAGATTGTAGCT
NR_144696.1      TGCCTCATAGAGGGGAATAGCCTTCCGAAAGGAAGATTAATACCGCATAAGATTGTAGCT
ettb2            TGCCTCATAGAGGGGAATAGCCTTTCGAAAGGAAGATTAATACCGCATAAGATTGTAGCT
                 ************************ ***********************************

ettb3            TCGCATGAAGTAGCAATTAAAGGAGCAATCCGCTATGAGATGGGCCCGCGGCGCATTAGC
ettb1            TCGCATGAAGTAGCAATTAAAGGAGCAATCCGCTATGAGATGGGCCCGCGGCGCATTAGC
NR_144696.1      TCGCATGAAGTAGCAATTAAAGGAGCAATCCGCTATGAGATGGGCCCGCGGCGCATTAGC
ettb2            TCGCATGAAGTAGCAATTAAAGGAGCAATCCGCTATGAGATGGGCCCGCGGCGCATTAGC
                 ************************************************************

ettb3            TAGTTGGTGAGGTAACGGCTCACCAAGGCGACGATGCGTAGCCGACCTGAGAGGGTGATC
ettb1            TAGTTGGTGAGGTAACGGCTCACCAAGGCGACGATGCGTAGCCGACCTGAGAGGGTGATC
NR_144696.1      TAGTTGGTGAGGTAACGGCTCACCAAGGCGACGATGCGTAGCCGACCTGAGAGGGTGATC
ettb2            TAGTTGGTGAGGTAACGGCTCACCAAGGCGACGATGCGTAGCCGACCTGAGAGGGTGATC
```

```
                    ************************************************************

ettb3               GGCCACATTGGGACTGAGACACGGCCCAGACTCCTACGGGAGGCAGCAGTGGGGAATATT
ettb1               GGCCACATTGGGACTGAGACACGGCCCAGACTCCTACGGGAGGCAGCAGTGGGGAATATT
NR_144696.1         GGCCACATTGGGACTGAGACACGGCCCAGACTCCTACGGGAGGCAGCAGTGGGGAATATT
ettb2               GGCCACATTGGGACTGAGACACGGCCCAGACTCCTACGGGAGGCAGCAGTGGGGAATATT
                    ************************************************************

ettb3               GCACAATGGGGGAAACCCTGAT-GCAGCAACGCCGCGTGAGTGAAGACGGCCTTCGGGGT
ettb1               GCACAATGGGGGAAACCCTGATTGCAGCAACCCCGCGTGAGTGATGACGGCCTTCGGG-T
NR_144696.1         GCACAATGGGGGAAACCCTGAT-GCAGCAACGCCGCGTGAGTGATGACGGCCTTCGGG-T
ettb2               GCACAATGGGGGAAACCCTGAGGCAGACAACCCCGCGTGAATGATGACGGCCTTCGGGGT
                    *********************      **** ******** *** ************* *

ettb3               TGAAAA-GCTCTGTCTTCAGGGACGATAATGACGGTACCTGGAGGAGGAAGCCACGGCAA
ettb1               TGTAAA-GCTCTGTCTTCAGGGACGATAATGACGGTACCTG-AGGAGGAAGCCACGGCTA
NR_144696.1         TGTAAA-GCTCTGTCTTCAGGGACGATAATGACGGTACCTG-AGGAGGAAGCCACGGCTA
ettb2               TGTAAAAGCTCTGTCTTCAGGGAAGATAATGACGGTACCTG-AGGAGGAAGCCACGGCAA
                    ** *** **************** ***************** **************** *

ettb3               ACTACGTGCCAGCAGCCGCGGTAATACGTAGGTGGCAAGCGTTGTCCGGATTTACTGGGC
ettb1               ACTACGTGCCAGCAGCCGCGGTAATACGTAGGTGGCGAGCGTTGTCCGGATTTACTGGGC
NR_144696.1         ACTACGTGCCAGCAGCCGCGGTAATACGTAGGTGGCGAGCGTTGTCCGGATTTACTGGGC
ettb2               ACTACGTGCCAGCAGCCGCGGTAATACGTAGGTGGCGAGCGTTGTCCGGATTTACTGGGC
                    ************************************ ***********************

ettb3               GTAAAGGGAGCGTAGGCGGACTTTTAAGTGAGATGTGAAATACCCGGGCTCAACTTGGGT
ettb1               GTAAAGGGAGCGTAGGCGGACTTTTAAGTGAGATGTGAAATACCCGGGCTCAACTTGGGT
NR_144696.1         GTAAAGGGAGCGTAGGCGGACTTTTAAGTGAGATGTGAAATACCCGGGCTCAACTTGGGT
ettb2               GTAAAGGGAGCGTAGGCGGACTTTTAAGTGAGATGTGAAATACCCGGGCTCAACTTGGGT
                    ************************************************************

ettb3               GCTGCATTTCAAACTGGAAGTCTAGAGTGCAGGAGAGGAGAATGGAATTCCTAGTGTAGC
ettb1               GCTGCATTTCAAACTGGAAGTCTAGAGTGCAGGAGAGGAGAATGGAATTCCTAGTGTAGC
NR_144696.1         GCTGCATTTCAAACTGGAAGTCTAGAGTGCAGGAGAGGAGAATGGAATTCCTAGTGTAGC
ettb2               GCTGCATTTCAAACTGGAAGTCTAGAGTGCAGGAGAGGAGAATGGAATTCCTAGTGTAGC
                    ************************************************************

ettb3               GGTGAAATGCGTAGAGATTAGGAAGAACACCAGTGGCGAAGGCGATTCTCTGGACTGTAA
ettb1               GGTGAAATGCGTAGAGATTAGGAAGAACACCAGTGGCGAAGGCGATTCTCTGGACTGTAA
NR_144696.1         GGTGAAATGCGTAGAGATTAGGAAGAACACCAGTGGCGAAGGCGATTCTCTGGACTGTAA
ettb2               GGTGAAATGCGTAGAGATTAGGAAGAACACCAGTGGCGAAGGCGATTCTCTGGACTGTAA
                    ************************************************************

ettb3               CTGACGCTGAGGCTCGAAAGCGTGGGGAGCAAACAGGATTAGATACCCTGGTAGTCCACG
ettb1               CTGACGCTGAGGCTCGAAAGCGTGGGGAGCAAACAGGATTAGATACCCTGGTAGTCCACG
NR_144696.1         CTGACGCTGAGGCTCGAAAGCGTGGGGAGCAAACAGGATTAGATACCCTGGTAGTCCACG
ettb2               CTGACGCTGAGGCTCGAAAGCGTGGGGAGCAAACAGGATTAGATACCCTGGTAGTCCACG
                    ************************************************************

ettb3               CCGTAAACGATGAATACTAGGTGGTAGGGGTTGTCATGACCTCTGTGCCGCCGCTAACGC
ettb1               CCGTAAACGATGAATACTAGGTG-TAGGGGTTGTCATGACCTCTGTGCCGCCGCTAACGC
NR_144696.1         CCGTAAACGATGAATACTAGGTG-TAGGGGTTGTCATGACCTCTGTGCCGCCGCTAACGC
ettb2               CCGTAAACGATGAATACTAGGTG-TAGGGGTTGTCATGACCTCTGTGCCGCCGCTAACGC
                    *********************** ************************************

ettb3               ATTAAGTATTCCGCCTGGGGAGTACGGTCGCAAGATTAAAACTCAAAGGAATTGACGGGG
ettb1               ATTAAGTATTCCGCCTGGGGAGTACGGTCGCAAGATTAAAACTCAAAGGAATTGACGGGG
NR_144696.1         ATTAAGTATTCCGCCTGGGGAGTACGGTCGCAAGATTAAAACTCAAAGGAATTGACGGGG
ettb2               ATTAAGTATTCCGCCTGGGGAGTACGGTCGCAAGATTAAAACTCAAAGGAATTGACGGGG
                    ************************************************************

ettb3               GCCCGCACAAGCAGCGGAGCATGTGGTTTAATTCGAAGCAACGCGAAGAACCTTACCTAG
ettb1               GCCCGCACAAGCAGCGGAGCATGTGGTTTAATTCGAAGCAACGCGAAGAACCTTACCTAG
NR_144696.1         GCCCGCACAAGCAGCGGAGCATGTGGTTTAATTCGAAGCAACGCGAAGAACCTTACCTAG
ettb2               GCCCGCACAAGCAGCGGAGCATGTGGTTTAATTCGAAGCAACGCGAAGAACCTTACCTAG
                    ************************************************************
```

```
ettb3           ACTTGACATCTCCTGCATTACCCTTAATCGGGAAAGTTCCTTCGGGGACAGGAAGACAGG
ettb1           ACTTGACATCTCCTGCATTACCCTTAATCGGGAAAGTTCCTTCGGGGACAGGAAGACAGG
NR_144696.1     ACTTGACATCTCCTGCATTACCCTTAATCGGGGAAGTTCCTTCGGGAACAGGAAGACAGG
ettb2           ACTTGACATCTCCTGCATTACCCTTAATCGGGAAAGTTCCTTCGGGGACAGGAAGACAGG
                ******************************** ************* *************

ettb3           TGGTGCATGGTTGTCGTCAGCTCC-TGTCCTG-AGAAGTTTGGGTTAAATCCCGCAACGA
ettb1           TGGTGCATGGTTGTCGTCAGCTCC-TGTCCTG-AGATGTTTGGGTTAAGTCCCGCAACGA
NR_144696.1     TGGTGCATGGTTGTCGTCAGCTCG-TGTCGTG-AGATGTT-GGGTTAAGTCCCGCAACGA
ettb2           TGGTGCATGGTTGTCCTCAACTCCGTGTCCGGGAAATGTTGGGGTTAAGTCCCCCAACAA
                *************** *** ***  ****  * * * *** ******* **** **** *

ettb3           GCGCAACCCTTATTGTTAGTTGCTACC-ATTTAGTTGAGCACTCTAGCGAGACTGCCCGG
ettb1           GCGCAACCCTTATTGGTAATTGCTACCCATTTAGTTGAGCACTCTAGCGAGACTGCCCGG
NR_144696.1     GCGCAACCCTTATTGTTAGTTGCTACC-ATTTAGTTGAGCACTCTAGCGAGACTGCCCGG
ettb2           GCGCAACCCCTATTGGTAGTGGCCACC-ATTTAGTTGAGAACTCTAGCGAGACTGCCCGG
                ********* ***** ** * ** *** *********** ********************

ettb3           GTTAACCGGGAGGAAGGTGGGGATGACGTCAAATCATCATGCCCCTTATGTCTAGGGCTA
ettb1           GTTAACCGGGAGGAAGGTGGGGATGACGTCAAATCATCATGCCCCTTATGTCTAGGGCTA
NR_144696.1     GTTAACCGGGAGGAAGGTGGGGATGACGTCAAATCATCATGCCCCTTATGTCTAGGGCTA
ettb2           GTTAACCGGGAGGAAGGTGGGGATGACGTCAAATCATCATGCCCCTTATGTCTAGGGCTA
                ************************************************************

ettb3           CACACGTGCTACAATGGCAAGTACAAAGAGACGCAATACCGCGAGGTGGAGCAAAACTCA
ettb1           CACACGTGCTACAATGGCAAGTACAAAGAGACGCAATACCGCGAGGTGGAGCAAAACTCA
NR_144696.1     CACACGTGCTACAATGGCAAGTACAAAGAGAAGCAAGACCGCGAGGTGGAGCAAAACTCA
ettb2           CACACGTGCTACAATGGCAAGTACAAAGAGACGCAATACCGCGAGGTGGAGCAAAACTCA
                ******************************* **** ***********************

ettb3           AAAACTTGTCTCAGTTCGGATTGTAGGCTGAAACTCGCCTACATGAAGCTGGAGTTGCTA
ettb1           AAAACTTGTCTCAGTTCGGATTGTAGGCTGAAACTCGCCTACATGAAGCTGGAGTTGCTA
NR_144696.1     AAAACTTGTCTCAGTTCGGATTGTAGGCTGAAACTCGCCTACATGAAGCTGGAGTTGCTA
ettb2           AAAACTTGTCTCAGTTCGGATTGTAGGCTGAAACTCGCCTACATGAAGCTGGAGTTGCTA
                ************************************************************

ettb3           GTAATCGCGAATCAGCATGTCGCGGTGAATACGTTCCCGGGCCTTGTACACACCGCCCGT
ettb1           GTAATCGCGAATCAGCATGTCGCGGTGAATACGTTCCCGGGCCTTGTACACACCGCCCGT
NR_144696.1     GTAATCGCGAATCAGCATGTCGCGGTGAATACGTTCCCGGGCCTTGTACACACCGCCCGT
ettb2           GTAATCGCGAATCAGCATGTCGCGGTGAATACGTTCCCGGGCCTTGTACACACCGCCCGT
                ************************************************************

ettb3           CACACCATGAGAGTTGGCAATACCCAAAGTGCGTGATCTAACTCGCAAGAGAGGAAGCGC
ettb1           CACACCATGAGAGTTGGCAATACCCAAAGTGCGTGATCTAACTCGCAAGAGAGGAAGCGC
NR_144696.1     CACACCATGAGAGTTGGCAATACCCAAAGTGCGTGATCTGACTCGCAAGAGAGGAAGCGC
ettb2           CACACCATGAGAGTTGGCAATACCCAAAGTGCGTGATCTAACTCGCAAGAGAGGAAGCGC
                *************************************** ********************

ettb3           CCTAAG--GTAGATCCGGA-----------------------------
ettb1           CCTAAGTAGAAAGTCT--------------------------------
NR_144696.1     CCTAAG-GTAGGGTCAGCGATTGGGTGAAGTCGTAACAACGTAGCCGT
ettb2           CCTAAG---GTAGTCAGGCA----------------------------
                ******       **
```

clustalw.dnd

```
(
NR_144696.1:0.01319,
(
ettb3:0.00756,
ettb1:0.00939)
:0.00197,
ettb2:0.01494);
```

16S rDNA sequences used for the analysis:

>*Clostridium filamentum* ettb1

CGTACGCTGCTAACCTGCAGTCGAGCGAGTGAATCTCCTTCGGGAGTGAAGCTAGCGGCGGACGGGTGAGTA
ACACGTGGGCAACCTGCCTCATAGAGGGGAATAGCCTTTCGAAAGGAAGATTAATACCGCATAAGATTGTAGC
TTCGCATGAAGTAGCAATTAAAGGAGCAATCCGCTATGAGATGGGCCCGCGGCGCATTAGCTAGTTGGTGAG
GTAACGGCTCACCAAGGCGACGATGCGTAGCCGACCTGAGAGGGTGATCGGCCACATTGGGACTGAGACACG
GCCCAGACTCCTACGGGAGGCAGCAGTGGGGAATATTGCACAATGGGGGAAACCCTGATTGCAGCAACCCCG
CGTGAGTGATGACGGCCTTCGGGTTGTAAAGCTCTGTCTTCAGGGACGATAATGACGGTACCTGAGGAGGAA
GCCACGGCTAACTACGTGCCAGCAGCCGCGGTAATACGTAGGTGGCGAGCGTTGTCCGGATTTACTGGGCGT
AAAGGGAGCGTAGGCGGACTTTTAAGTGAGATGTGAAATACCCGGGCTCAACTTGGGTGCTGCATTTCAAACT
GGAAGTCTAGAGTGCAGGAGAGGAGAATGGAATTCCTAGTGTAGCGGTGAAATGCGTAGAGATTAGGAAGA
ACACCAGTGGCGAAGGCGATTCTCTGGACTGTAACTGACGCTGAGGCTCGAAAGCGTGGGGAGCAAACAGGA
TTAGATACCCTGGTAGTCCACGCCGTAAACGATGAATACTAGGTGTAGGGGTTGTCATGACCTCTGTGCCGCC
GCTAACGCATTAAGTATTCCGCCTGGGGAGTACGGTCGCAAGATTAAAACTCAAAGGAATTGACGGGGGCCC
GCACAAGCAGCGGAGCATGTGGTTTAATTCGAAGCAACGCGAAGAACCTTACCTAGACTTGACATCTCCTGCA
TTACCCTTAATCGGGAAAGTTCCTTCGGGGACAGGAAGACAGGTGGTGCATGGTTGTCGTCAGCTCCTGTCCT
GAGATGTTTGGGTTAAGTCCCGCAACGAGCGCAACCCTTATTGGTAATTGCTACCCATTTAGTTGAGCACTCTA
GCGAGACTGCCCGGGTTAACCGGGAGGAAGGTGGGGATGACGTCAAATCATCATGCCCCTTATGTCTAGGGC
TACACACGTGCTACAATGGCAAGTACAAAGAGACGCAATACCGCGAGGTGGAGCAAAACTCAAAAACTTGTCT
CAGTTCGGATTGTAGGCTGAAACTCGCCTACATGAAGCTGGAGTTGCTAGTAATCGCGAATCAGCATGTCGCG
GTGAATACGTTCCCGGGCCTTGTACACACCGCCCGTCACACCATGAGAGTTGGCAATACCCAAAGTGCGTGAT
CTAACTCGCAAGAGAGGAAGCGCCCTAAGTAGAAAGTCT

>*Clostridium filamentum* ettb2

CAAGGCGGCTGCTAAAATGCAGTCGAGCGAGTGAATCTCCTTCGGGAGTGAAGCTAGCGGCGGACGGGTGA
GTAACACGTGGGCAACCTGCCTCATAGAGGGGAATAGCCTTTCGAAAGGAAGATTAATACCGCATAAGATTGT
AGCTTCGCATGAAGTAGCAATTAAAGGAGCAATCCGCTATGAGATGGGCCCGCGGCGCATTAGCTAGTTGGT
GAGGTAACGGCTCACCAAGGCGACGATGCGTAGCCGACCTGAGAGGGTGATCGGCCACATTGGGACTGAGA
CACGGCCCAGACTCCTACGGGAGGCAGCAGTGGGGAATATTGCACAATGGGGGAAACCCTGAGGCAGACAA
CCCCGCGTGAATGATGACGGCCTTCGGGGTTGTAAAAGCTCTGTCTTCAGGGAAGATAATGACGGTACCTGAG
GAGGAAGCCACGGCAAACTACGTGCCAGCAGCCGCGGTAATACGTAGGTGGCGAGCGTTGTCCGGATTTACT
GGGCGTAAAGGGAGCGTAGGCGGACTTTTAAGTGAGATGTGAAATACCCGGGCTCAACTTGGGTGCTGCATT
TCAAACTGGAAGTCTAGAGTGCAGGAGAGGAGAATGGAATTCCTAGTGTAGCGGTGAAATGCGTAGAGATTA
GGAAGAACACCAGTGGCGAAGGCGATTCTCTGGACTGTAACTGACGCTGAGGCTCGAAAGCGTGGGGAGCA
AACAGGATTAGATACCCTGGTAGTCCACGCCGTAAACGATGAATACTAGGTGTAGGGGTTGTCATGACCTCTG
TGCCGCCGCTAACGCATTAAGTATTCCGCCTGGGGAGTACGGTCGCAAGATTAAAACTCAAAGGAATTGACGG
GGGCCCGCACAAGCAGCGGAGCATGTGGTTTAATTCGAAGCAACGCGAAGAACCTTACCTAGACTTGACATCT
CCTGCATTACCCTTAATCGGGAAAGTTCCTTCGGGGACAGGAAGACAGGTGGTGCATGGTTGTCCTCAACTCC
GTGTCCGGGAAATGTTGGGGTTAAGTCCCCCAACAAGCGCAACCCCTATTGGTAGTGGCCACCATTTAGTTGA
GAACTCTAGCGAGACTGCCCGGGTTAACCGGGAGGAAGGTGGGGATGACGTCAAATCATCATGCCCCTTATG
TCTAGGGCTACACACGTGCTACAATGGCAAGTACAAAGAGACGCAATACCGCGAGGTGGAGCAAAACTCAAA
AACTTGTCTCAGTTCGGATTGTAGGCTGAAACTCGCCTACATGAAGCTGGAGTTGCTAGTAATCGCGAATCAG
CATGTCGCGGTGAATACGTTCCCGGGCCTTGTACACACCGCCCGTCACACCATGAGAGTTGGCAATACCCAAA
GTGCGTGATCTAACTCGCAAGAGAGGAAGCGCCCTAAGGTAGTCAGGCA

>*Clostridium filamentum* ettb3

GGAACGGGGTGCTAACATGCAGTCGAGCGAGTGAATCTCCTTCGGGAGTGAAGCTAGCGGCGGACGGGTGA
GTAACACGTGGGCAACCTGCCTCATAGAGGGGAATAGCCTTTCGAAAGGAAGATTAATACCGCATAAGATTGT
AGCTTCGCATGAAGTAGCAATTAAAGGAGCAATCCGCTATGAGATGGGCCCGCGGCGCATTAGCTAGTTGGT
GAGGTAACGGCTCACCAAGGCGACGATGCGTAGCCGACCTGAGAGGGTGATCGGCCACATTGGGACTGAGA
CACGGCCCAGACTCCTACGGGAGGCAGCAGTGGGGAATATTGCACAATGGGGGAAACCCTGATGCAGCAACG
CCGCGTGAGTGAAGACGGCCTTCGGGGTTGAAAAGCTCTGTCTTCAGGGACGATAATGACGGTACCTGGAGG
AGGAAGCCACGGCAAACTACGTGCCAGCAGCCGCGGTAATACGTAGGTGGCAAGCGTTGTCCGGATTTACTG
GGCGTAAAGGGAGCGTAGGCGGACTTTTAAGTGAGATGTGAAATACCCGGGCTCAACTTGGGTGCTGCATTT
CAAACTGGAAGTCTAGAGTGCAGGAGAGGAGAATGGAATTCCTAGTGTAGCGGTGAAATGCGTAGAGATTAG
GAAGAACACCAGTGGCGAAGGCGATTCTCTGGACTGTAACTGACGCTGAGGCTCGAAAGCGTGGGGAGCAA
ACAGGATTAGATACCCTGGTAGTCCACGCCGTAAACGATGAATACTAGGTGGTAGGGGTTGTCATGACCTCTG
TGCCGCCGCTAACGCATTAAGTATTCCGCCTGGGGAGTACGGTCGCAAGATTAAAACTCAAAGGAATTGACGG
GGGCCCGCACAAGCAGCGGAGCATGTGGTTTAATTCGAAGCAACGCGAAGAACCTTACCTAGACTTGACATCT
CCTGCATTACCCTTAATCGGGAAAGTTCCTTCGGGGACAGGAAGACAGGTGGTGCATGGTTGTCGTCAGCTCC
TGTCCTGAGAAGTTTGGGTTAAATCCCGCAACGAGCGCAACCCTTATTGTTAGTTGCTACCATTTAGTTGAGCA
CTCTAGCGAGACTGCCCGGGTTAACCGGGAGGAAGGTGGGGATGACGTCAAATCATCATGCCCCTTATGTCTA
GGGCTACACACGTGCTACAATGGCAAGTACAAAGAGACGCAATACCGCGAGGTGGAGCAAAACTCAAAAACT
TGTCTCAGTTCGGATTGTAGGCTGAAACTCGCCTACATGAAGCTGGAGTTGCTAGTAATCGCGAATCAGCATG
TCGCGGTGAATACGTTCCCGGGCCTTGTACACACCGCCCGTCACACCATGAGAGTTGGCAATACCCAAAGTGC
GTGATCTAACTCGCAAGAGAGGAAGCGCCCTAAGGTAGATCCGGA

>NR_144696.1 *C saudiense* JCC

AGAGTTTGATCCTGGCTCAGGACGAACGCTGGCGGCGTGCCTAACACATGCAAGTCGAGCGAGTGAATCT

CCTTCGGGAGTGAAGCTAGCGGCGGACGGGTGAGTAACACGTGGGCAACCTGCCTCATAGAGGGGAATAG

CCTTCCGAAAGGAAGATTAATACCGCATAAGATTGTAGCTTCGCATGAAGTAGCAATTAAAGGAGCAATC

CGCTATGAGATGGGCCCGCGGCGCATTAGCTAGTTGGTGAGGTAACGGCTCACCAAGGCGACGATGCGTA

GCCGACCTGAGAGGGTGATCGGCCACATTGGGACTGAGACACGGCCCAGACTCCTACGGGAGGCAGCAGT

GGGGAATATTGCACAATGGGGGAAACCCTGATGCAGCAACGCCGCGTGAGTGATGACGGCCTTCGGGTTG

TAAAGCTCTGTCTTCAGGGACGATAATGACGGTACCTGAGGAGGAAGCCACGGCTAACTACGTGCCAGCA

GCCGCGGTAATACGTAGGTGGCGAGCGTTGTCCGGATTTACTGGGCGTAAAGGGAGCGTAGGCGGACTTT

TAAGTGAGATGTGAAATACCCGGGCTCAACTTGGGTGCTGCATTTCAAACTGGAAGTCTAGAGTGCAGGA

GAGGAGAATGGAATTCCTAGTGTAGCGGTGAAATGCGTAGAGATTAGGAAGAACACCAGTGGCGAAGGCG

ATTCTCTGGACTGTAACTGACGCTGAGGCTCGAAAGCGTGGGGAGCAAACAGGATTAGATACCCTGGTAG

TCCACGCCGTAAACGATGAATACTAGGTGTAGGGGTTGTCATGACCTCTGTGCCGCCGCTAACGCATTAA

GTATTCCGCCTGGGGAGTACGGTCGCAAGATTAAAACTCAAAGGAATTGACGGGGGCCCGCACAAGCAGC

GGAGCATGTGGTTTAATTCGAAGCAACGCGAAGAACCTTACCTAGACTTGACATCTCCTGCATTACCCTT

AATCGGGGAAGTTCCTTCGGGAACAGGAAGACAGGTGGTGCATGGTTGTCGTCAGCTCGTGTCGTGAGAT

GTTGGGTTAAGTCCCGCAACGAGCGCAACCCTTATTGTTAGTTGCTACCATTTAGTTGAGCACTCTAGCG

AGACTGCCCGGGTTAACCGGGAGGAAGGTGGGGATGACGTCAAATCATCATGCCCCTTATGTCTAGGGCT

ACACACGTGCTACAATGGCAAGTACAAAGAGAAGCAAGACCGCGAGGTGGAGCAAAACTCAAAAACTTGT

CTCAGTTCGGATTGTAGGCTGAAACTCGCCTACATGAAGCTGGAGTTGCTAGTAATCGCGAATCAGCATG

TCGCGGTGAATACGTTCCCGGGCCTTGTACACACCGCCCGTCACACCATGAGAGTTGGCAATACCCAAAG

TGCGTGATCTGACTCGCAAGAGAGGAAGCGCCCTAAGGTAGGGTCAGCGATTGGGTGAAGTCGTAACAAC

GTAGCCGT

**Supplementary information 2**

```
CLUSTAL O(1.2.4) multiple sequence alignment of fliC gene of C. filamentum
ETTB1, ETTB2 and ETTB3


GJHHNENP_00275
atgaggattaacacaaacttaaatgccatgaaggcaataaaaaactctaataaaaatgtt   60
NFHHKBOH_00275
atgaggattaacacaaacttaaatgccatgaaggcaataaaaaactctaataaaaatgtt   60
GMHBADHI_00275
atgaggattaacacaaacttaaatgccatgaaggcaataaaaaactctaataaaaatgtt   60

************************************************************

GJHHNENP_00275
tcttcatctggaaattcaatgaaaaaattaatttcagggttaagcattaataaagctgca   120
NFHHKBOH_00275
tcttcatctggaaattcaatgaaaaaattaatttcagggttaagcattaataaagctgca   120
GMHBADHI_00275
tcttcatctggaaattcaatgaaaaaattaatttcagggttaagcattaataaagctgca   120

************************************************************

GJHHNENP_00275
gatgatgcagctggattagctatatctgaaaagatgagaagtcaaataagagggttaaat   180
NFHHKBOH_00275
gatgatgcagctggattagctatatctgaaaagatgagaagtcaaataagagggttaaat   180
GMHBADHI_00275
gatgatgcagctggattagctatatctgaaaagatgagaagtcaaataagagggttaaat   180

************************************************************

GJHHNENP_00275
caagcagaacaaaatgcacaagatggtattttaatgcttcaaacagcagaagggactatt   240
NFHHKBOH_00275
caagcagaacaaaatgcacaagatggtattttaatgcttcaaacagcagaagggactatt   240
GMHBADHI_00275
caagcagaacaaaatgcacaagatggtattttaatgcttcaaacagcagaagggactatt   240

************************************************************

GJHHNENP_00275
gaggaagttggaaatatagttcaaagaatgagagaattatctgttcaatgtgcaaatgaa   300
NFHHKBOH_00275
gaggaagttggaaatatagttcaaagaatgagagaattatctgttcaatgtgcaaatgaa   300
GMHBADHI_00275
gaggaagttggaaatatagttcaaagaatgagagaattatctgttcaatgtgcaaatgaa   300

************************************************************

GJHHNENP_00275
agtaattctggagaagatagagaaaaaatagtaatagaattaaatgagttacatgaagaa   360
NFHHKBOH_00275
agtaattctggagaagatagagaaaaaatagtaatagaattaaatgagttacatgaagaa   360
GMHBADHI_00275
agtaattctggagaagatagagaaaaaatagtaatagaattaaatgagttacatgaagaa   360
```

************************************************************

GJHHNENP_00275
attgatagaatatcaaactcaactcaatttaatgaaaaagacttattaaatggacaaact 420
NFHHKBOH_00275
attgatagaatatcaaactcaactcaatttaatgaaaaagacttattaaatggacaaact 420
GMHBADHI_00275
attgatagaatatcaaactcaactcaatttaatgaaaaagacttattaaatggacaaact 420

************************************************************

GJHHNENP_00275
gtgataagaattgaaaagcaatatactattacaaaaggtgcagcaatgaatgatcaaatt 480
NFHHKBOH_00275
gtgataagaattgaaaagcaatatactattacaaaaggtgcagcaatgaatgatcaaatt 480
GMHBADHI_00275
gtgataagaattgaaaagcaatatactattacaaaaggtgcagcaatgaatgatcaaatt 480

************************************************************

GJHHNENP_00275
tctaaaatacaagttcaaaggggatttgattttagtgatttagaccttggaggattaggc 540
NFHHKBOH_00275
tctaaaatacaagttcaaaggggatttgattttagtgatttagaccttggaggattaggc 540
GMHBADHI_00275
tctaaaatacaagttcaaaggggatttgattttagtgatttagaccttggaggattaggc 540

************************************************************

GJHHNENP_00275
tttgtaagtcataaacaaggtgagattagttttgagcctggaaaaaatattaattataag 600
NFHHKBOH_00275
tttgtaagtcataaacaaggtgagattagttttgagcctggaaaaaatattaattataag 600
GMHBADHI_00275
tttgtaagtcataaacaaggtgagattagttttgagcctggaaaaaatattaattataag 600

************************************************************

GJHHNENP_00275
gtttctgtagataatatggttaatcaaaaagtacaacctggtgctacaataactataaat 660
NFHHKBOH_00275
gtttctgtagataatatggttaatcaaaaagtacaacctggtgctacaataactataaat 660
GMHBADHI_00275
gtttctgtagataatatggttaatcaaaaagtacaacctggtgctacaataactataaat 660

************************************************************

GJHHNENP_00275
gattcaaatggaaaatatattaaatttgctgttaaacaagaaatagatttaaatattgca 720
NFHHKBOH_00275
gattcaaatggaaaatatattaaatttgctgttaaacaagaaatagatttaaatattgca 720
GMHBADHI_00275
gattcaaatggaaaatatattaaatttgctgttaaacaagaaatagatttaaatattgca 720

************************************************************

GJHHNENP_00275
actcaagttaaaatgataaatgtagcacttaaagaaaaagaaataaagggaaaagaaatt 780
NFHHKBOH_00275
actcaagttaaaatgataaatgtagcacttaaagaaaaagaaataaagggaaaagaaatt 780
GMHBADHI_00275
actcaagttaaaatgataaatgtagcacttaaagaaaaagaaataaagggaaaagaaatt 780

************************************************************

GJHHNENP_00275
aagttgcaaattggagcaaattcagatagctctcaaacattaaaagttaaaatagataac 840
NFHHKBOH_00275
aagttgcaaattggagcaaattcagatagctctcaaacattaaaagttaaaatagataac 840
GMHBADHI_00275
aagttgcaaattggagcaaattcagatagctctcaaacattaaaagttaaaatagataac 840

************************************************************

GJHHNENP_00275
atgagcacagaatcattggaaataagcaaagctgatatatcaaatatgaaaaaagaaggc 900
NFHHKBOH_00275
atgagcacagaatcattggaaataagcaaagctgatatatcaaatatgaaaaaagaaggc 900
GMHBADHI_00275
atgagcacagaatcattggaaataagcaaagctgatatatcaaatatgaaaaaagaaggc 900

************************************************************

GJHHNENP_00275
agtaaaggaacagaagctgctacaaaaatgatagataatttagatgatgctttagaaaga 960
NFHHKBOH_00275
agtaaaggaacagaagctgctacaaaaatgatagataatttagatgatgctttagaaaga 960
GMHBADHI_00275
agtaaaggaacagaagctgctacaaaaatgatagataatttagatgatgctttagaaaga 960

************************************************************

GJHHNENP_00275
gttaatgtttcaagggggaaccttggtgcaatgcaaaatagattgcaaacagcttcatct 1020
NFHHKBOH_00275
gttaatgtttcaagggggaaccttggtgcaatgcaaaatagattgcaaacagcttcatct 1020
GMHBADHI_00275
gttaatgtttcaagggggaaccttggtgcaatgcaaaatagattgcaaacagcttcatct 1020

************************************************************

GJHHNENP_00275
aacttaaatactatgaatgaaaacttaacttcaatagaatcaagaattagagatgtagat 1080
NFHHKBOH_00275
aacttaaatactatgaatgaaaacttaacttcaatagaatcaagaattagagatgtagat 1080
GMHBADHI_00275
aacttaaatactatgaatgaaaacttaacttcaatagaatcaagaattagagatgtagat 1080

************************************************************

```
GJHHNENP_00275
gttgcaaaagaaataatgaatttatcaaaaataaatttaataattcaagcaaatcaagca    1140
NFHHKBOH_00275
gttgcaaaagaaataatgaatttatcaaaaataaatttaataattcaagcaaatcaagca    1140
GMHBADHI_00275
gttgcaaaagaaataatgaatttatcaaaaataaatttaataattcaagcaaatcaagca    1140

************************************************************

GJHHNENP_00275      ataatggtgcaagcaaaatctcaaccagaagctgtaatacagttattaaaataa
        1194
NFHHKBOH_00275      ataatggtgcaagcaaaatctcaaccagaagctgtaatacagttattaaaataa
        1194
GMHBADHI_00275      ataatggtgcaagcaaaatctcaaccagaagctgtaatacagttattaaaataa
        1194
                    ******************************************************
```

Sequences used :

fliC sequence found in *C. filamentum* ETTB1

>GMHBADHI_00275 fliC A-type flagellin 270911:272104 forward

atgaggattaacacaaacttaaatgccatgaaggcaataaaaaactctaataaaaatgtt

tcttcatctggaaattcaatgaaaaaattaatttcagggttaagcattaataaagctgca

gatgatgcagctggattagctatatctgaaaagatgagaagtcaaataagagggttaaat

caagcagaacaaaatgcacaagatggtattttaatgcttcaaacagcagaagggactatt

gaggaagttggaaatatagttcaaagaatgagagaattatctgttcaatgtgcaaatgaa

agtaattctggagaagatagagaaaaaatagtaatagaattaaatgagttacatgaagaa

attgatagaatatcaaactcaactcaatttaatgaaaaagacttattaaatggacaaact

gtgataagaattgaaaagcaatatactattacaaaaggtgcagcaatgaatgatcaaatt

tctaaaatacaagttcaaaggggatttgattttagtgatttagaccttggaggattaggc

tttgtaagtcataaacaaggtgagattagttttgagcctggaaaaaaatattaattataag

gtttctgtagataatatggttaatcaaaaagtacaacctggtgctacaataactataaat

gattcaaatggaaaatatattaaatttgctgttaaacaagaaatagatttaaatattgca

actcaagttaaaatgataaatgtagcacttaaagaaaaagaaataaagggaaaagaaatt

aagttgcaaattggagcaaattcagatagctctcaaacattaaaagttaaaatagataac

atgagcacagaatcattggaaataagcaaagctgatatatcaaatatgaaaaaagaaggc

agtaaaggaacagaagctgctacaaaaatgatagataatttagatgatgctttagaaaga

gttaatgtttcaagggggaaccttggtgcaatgcaaaatagattgcaaacagcttcatct aacttaaatactatgaatgaaaacttaacttcaatagaatcaagaattagagatgtagat

gttgcaaaagaaataatgaatttatcaaaaataaatttaataattcaagcaaatcaagca

ataatggtgcaagcaaaatctcaaccagaagctgtaatacagttattaaaataa

fliC sequence found in *C. filamentum* ETTB2

>NFHHKBOH_00275 fliC A-type flagellin 270812:272005 forward

atgaggattaacacaaacttaaatgccatgaaggcaataaaaaactctaataaaaatgtt

tcttcatctggaaattcaatgaaaaaattaatttcagggttaagcattaataaagctgca

gatgatgcagctggattagctatatctgaaaagatgagaagtcaaataagagggttaaat

caagcagaacaaaatgcacaagatggtattttaatgcttcaaacagcagaagggactatt

gaggaagttggaaatatagttcaaagaatgagagaattatctgttcaatgtgcaaatgaa

agtaattctggagaagatagagaaaaaatagtaatagaattaaatgagttacatgaagaa

attgatagaatatcaaactcaactcaatttaatgaaaaagacttattaaatggacaaact

gtgataagaattgaaaagcaatatactattacaaaaggtgcagcaatgaatgatcaaatt

tctaaaatacaagttcaaaggggatttgattttagtgatttagaccttggaggattaggc

tttgtaagtcataaacaaggtgagattagttttgagcctggaaaaaatattaattataag

gtttctgtagataatatggttaatcaaaaagtacaacctggtgctacaataactataaat

gattcaaatggaaaatatattaaatttgctgttaaacaagaaatagatttaaatattgca

actcaagttaaaatgataaatgtagcacttaaagaaaaagaaataaagggaaaagaaatt

aagttgcaaattggagcaaattcagatagctctcaaacattaaaagttaaaatagataac

atgagcacagaatcattggaaataagcaaagctgatatatcaaatatgaaaaaagaaggc

agtaaaggaacagaagctgctacaaaaatgatagataatttagatgatgctttagaaaga

gttaatgtttcaagggggaaccttggtgcaatgcaaaatagattgcaaacagcttcatct

aacttaaatactatgaatgaaaacttaacttcaatagaatcaagaattagagatgtagat

gttgcaaaagaaataatgaatttatcaaaaataaatttaataattcaagcaaatcaagca

ataatggtgcaagcaaaatctcaaccagaagctgtaatacagttattaaaataa

fliC sequence found in *C. filamentum* ETTB3

>GJHHNENP_00275 fliC A-type flagellin 270935:272128 forward

atgaggattaacacaaacttaaatgccatgaaggcaataaaaaactctaataaaaatgtt

tcttcatctggaaattcaatgaaaaaattaatttcagggttaagcattaataaagctgca gatgatgcagctggattagctatatctgaaaagatgagaagtcaaataagagggttaaat

caagcagaacaaaatgcacaagatggtattttaatgcttcaaacagcagaagggactatt

gaggaagttggaaatatagttcaaagaatgagagaattatctgttcaatgtgcaaatgaa

agtaattctggagaagatagagaaaaaatagtaatagaattaaatgagttacatgaagaa

attgatagaatatcaaactcaactcaatttaatgaaaaagacttattaaatggacaaact

gtgataagaattgaaaagcaatatactattacaaaaggtgcagcaatgaatgatcaaatt

tctaaaatacaagttcaaaggggatttgattttagtgatttagaccttggaggattaggc

tttgtaagtcataaacaaggtgagattagttttgagcctggaaaaaatattaattataag

gtttctgtagataatatggttaatcaaaaagtacaacctggtgctacaataactataaat

gattcaaatggaaaatatattaaatttgctgttaaacaagaaatagatttaaatattgca

actcaagttaaaatgataaatgtagcacttaaagaaaaagaaataaagggaaaagaaatt

aagttgcaaattggagcaaattcagatagctctcaaacattaaaagttaaaatagataac

atgagcacagaatcattggaaataagcaaagctgatatatcaaatatgaaaaaagaaggc

agtaaaggaacagaagctgctacaaaaatgatagataatttagatgatgctttagaaaga

gttaatgtttcaagggggaaccttggtgcaatgcaaaatagattgcaaacagcttcatct

aacttaaatactatgaatgaaaacttaacttcaatagaatcaagaattagagatgtagat

gttgcaaaagaaataatgaatttatcaaaaataaatttaataattcaagcaaatcaagca

ataatggtgcaagcaaaatctcaaccagaagctgtaatacagttattaaaataa
